# RNA polymerase inhibitors reveal active-site motions essential for the nucleotide-addition cycle

**DOI:** 10.64898/2026.04.06.716786

**Authors:** Yukti Dhingra, Robert Landick, Elizabeth A. Campbell, Seth A. Darst

## Abstract

The nucleotide-addition cycle (NAC) of multi-subunit DNA-dependent RNA polymerases (RNAPs) involves coordinated conformational changes in conserved active-site structural elements, including the trigger loop (TL). The TL is open (unfolded) in most RNAP structures but can close (fold) in substrate-bound (post- or pre-translocated) states of the RNAP, promoting catalysis. TL closure has been associated with closure of another conserved structural element, the Rim-Helices/F-loop (RH-FL), but the role of the RH-FL in the NAC is unclear. Antibiotic leads CBR9379 and AAP-SO_2_ inhibit the *Escherichia coli* and *Mycobacterium tuberculosis* RNAPs, respectively, by binding in a pocket formed by the bridge helix and RH-FL. The precise mechanism of action for these inhibitors is yet to be defined. We present cryo-electron microscopy structures showing that both compounds inhibit the RNAP NAC by preventing RH-FL closure, thereby allosterically destabilizing the closed TL. This work reveals a conserved mechanistic principle of RNAP catalysis across all domains of life and provides new insight for antibiotic design.

## INTRODUCTION

Cellular RNA polymerases (RNAPs) are multi-subunit, highly processive enzymes that catalyze RNA synthesis in a DNA-dependent manner, adding ribonucleotides to a growing RNA chain through a nucleotide-addition cycle (NAC; Fig. 1A). The NAC comprises a sequence of molecular events (1, 2) accompanied by coordinated conformational changes in conserved structural elements of the RNAP, most prominently the trigger loop (TL). The TL is an unstructured loop (open) in most RNAP structures but folds into a helical hairpin (closed) when a cognate NTP is bound in the active site (3–7) (Fig. 1A-C).

**Figure 1.**
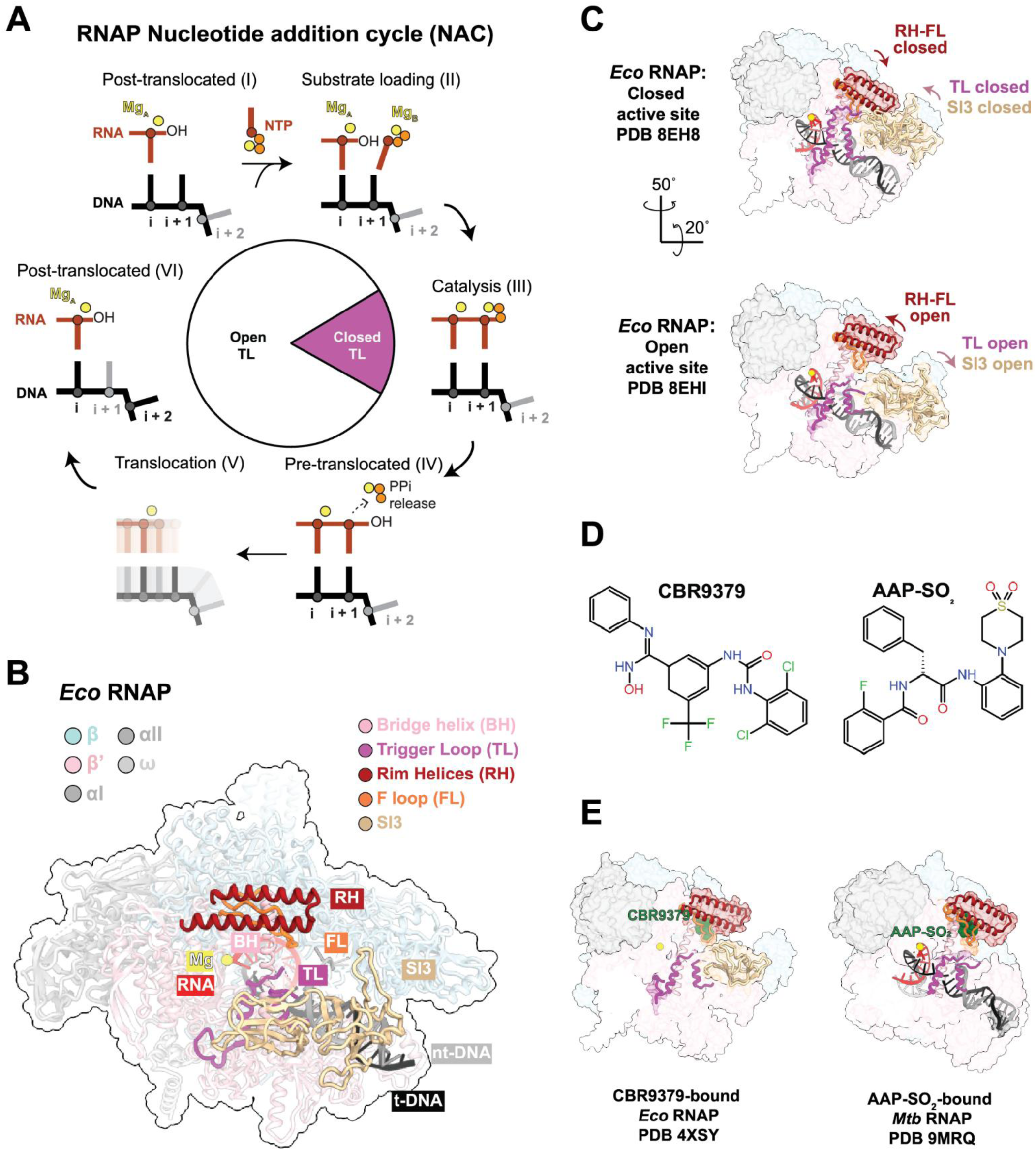
RNAP nucleotide-addition cycle, conserved structural elements of the RNAP active site, and allosteric inhibitors of the *Eco* and *Mtb* RNAP NAC. (A) Schematic illustration of the RNAP nucleotide addition cycle. The cycle begins in the post-translocated state (I), with an empty i+1 site and an open TL that permits entry of a nucleoside triphosphate (NTP) substrate into the active site. During substrate loading (II), the incoming NTP, coordinated by a Mg²⁺ ion, base pairs with the complementary t-strand DNA base at the i+1 site. Correct Watson–Crick pairing induces TL closing (folding). Catalysis (III) results in the formation of a phosphodiester bond between the 3′-oxygen of the RNA transcript and the α-phosphate of the NTP, followed by TL opening and PPi release (IV). The nucleic acids in the resulting pre-translocated state (IV) translocate one base pair with respect to the RNAP (V), giving rise to a new post-translocated state (VI), where the cycle begins again. (B) Structure of *Eco*RNAP (PDB 8EHI), shown in cartoon and colored by RNAP subunit but highlighting key structural motifs (color-key shown). RNA is colored red, t-strand DNA is black, nt-strand DNA is dark gray. The RNAP active-site Mg^2+^ is yellow. (C) Surface view of *Eco*RNAP active-site closed (top) and open (bottom) structures, color-coded as in (B). Arrows indicate closed and open RH-FL, TL and SI3. (C) Chemical structures of allosteric RNAP inhibitors CBR9379 and AAP-SO_2_. (D) Left: Surface view of CBR9379-bound *Eco*RNAP. Right: Surface view of AAP-SO_2_-bound *Mtb*EC. Color-coding as in (B), with the antibiotics colored green.

The NAC starts from a post-translocated, TL-open state [Fig. 1A(I)] permitting Mg²⁺-coordinated NTP binding and base pairing at the i+1 site (II), followed by TL closure, triggering catalysis (III) through direct contacts with the NTP substrate (3, 5, 8, 9). Phosphodiester bond formation is followed by TL opening and pyrophosphate (PPi) release in a pre-translocated state (IV), and translocation (V) by one base pair to regenerate a post-translocated state (VI) (10, 11).

Another conserved RNAP structural element, the Rim-Helices/F-loop (RH-FL; Fig. 1B) has sometimes (but not always) been observed to close toward the active site when the TL closes (12–14) (Fig. 1C). The FL strongly influences the NAC; swapping FL sequences between bacterial RNAPs can confer elongation behaviors characteristic of the donor species (15, 16). Compared with the bridge helix (BH) and TL, FL sequences are more diverse across bacteria (Fig. S1). RH-FL closure may be coordinated with TL closure since a closed RH-FL has not been observed without a closed TL, but the structural role of the RH-FL in the NAC is unclear. Each of these structural elements crucial for the NAC (RH-FL, BH, TL) are conserved in RNAP structures from bacteria to humans (Fig. S2A-D).

In some bacteria, such as *Escherichia coli* (*Eco*), the TL contains a large (188 residue) sequence insertion, SI3 (17, 18), which undergoes coordinated movements with the TL (13, 19). When the TL closes, SI3 also closes toward the active site, and when the TL opens, SI3 opens (Fig. 1C).

Because progression through the NAC is closely coordinated with RNAP conformational changes, small-molecule or peptidic inhibitors can bind relatively far from the RNAP active site Mg^2+^-ion (i.e. > 28 Å) but perturb the NAC allosterically by interfering with active-site motions (20–25). The CBR series of compounds (Fig. 1D) were identified as allosteric inhibitors of *Eco*RNAP that perturb the NAC by inhibiting RNAP catalytic activities (but not NTP substrate binding and not translocation) via contacts with the FL (26). Subsequent X-ray crystal structures delineated atomic details of the CBR binding poses and CBR–RNAP interactions (Fig. 1E; Fig. S1) (27, 28). The CBRs bind in a surface-exposed pocket, contacting the BH, the FL, and nearby parts of the β-subunit (Fig. 1E). The CBRs inhibit catalysis by perturbing TL closing but do not directly contact the TL (Fig. S1) (27). Collectively, studies of the CBR inhibition mechanism have led to the inference that the CBRs allosterically interfere with RNAP active-site motions important for catalysis (26, 27), but precisely what RNAP motions are affected by CBR binding, and how, is not clear.

The CBRs selectively inhibit Gram-negative bacteria and have activity against some Gram-positive bacteria (29, 30). *In vitro*, the CBRs do not inhibit human RNAP II (Pol II) nor RNAP from the major human pathogen *Mycobacterium tuberculosis* (*Mtb*) due to sequence differences in the binding pocket (Fig. S1) (27). However, *N*a-aroyl-*N*-aryl-phenylalanineamides, such as AAP-SO_2_ (Fig. 1D) (31) were found to inhibit *Mtb*RNAP and bind in the same respective pocket (Fig. 1E), leading to the inference that the inhibition mechanism for the CBRs and AAPs, while unknown in detail, is the same (32, 33).These properties make CBRs and AAPs promising leads for antibiotic development (33).

In this work, we determined a series of cryo-electron microscopy (cryo-EM) structures of *Eco*RNAP elongation complexes (ECs) ± CBR9379, and *Mtb*RNAP ECs ± AAP-SO_2_. Analysis of the structures establishes the mechanism of action of CBR and AAP inhibitors. CBR9379 and AAP-SO_2_ prevent closure of the active site for their respective RNAPs by holding the RH-FL domain in the open position. We further conclude that RH-FL closing in coordination with TL closing is crucial for efficient progression through the NAC. We propose that this mechanism is conserved across all cellular RNAPs due to the preserved architecture of these structural elements (Fig. S2A-D) (34).

## RESULTS

### CBR9379 dramatically slows the *Eco*RNAP single-nucleotide addition cycle

Comparison of *Eco*RNAP X-ray crystal structures with and without bound CBRs did not reveal RNAP conformation changes that explain the mechanism of CBR inhibition (27, 28). Moreover, crystal packing effects can complicate the interpretation of conformational changes observed in crystal structures. We reasoned that single particle cryo-EM would allow us to examine shifts in RNAP conformational ensembles induced by CBR binding, and help elucidate the structural mechanism for inhibition, information inaccessible to X-ray crystallography (13). We previously used cryo-EM to examine *Eco*RNAP on an elemental pause sequence (the *Eco his* operon leader sequence with the RNA pause hairpin deleted) (34–36) in an elemental paused EC (*his*-ePEC) and observed many distinct conformational states (13). Additionally, CBRs have previously been shown to preferentially inhibit nucleotide addition at the *his* pause (26). Therefore, we sought to examine the effect of CBR9379 (Fig. 1D) on the *Eco*RNAP *his*-ePEC conformational ensemble.

To confirm the effect of CBR9379 on the nucleic-acid scaffold used for cryo-EM (Fig. 2A), we assembled an *Eco*RNAP *his*-ePEC using an 18-nt RNA with a C at the 3′ end (C18-RNA) and extended it to the U19 pause site using [a-^32^P]UTP (Fig. 2A-B). GTP (10 mM) was then added, and single-nucleotide extension from U19 to G20 was monitored as a function of time (Fig. 2B-C).

**Figure 2.**
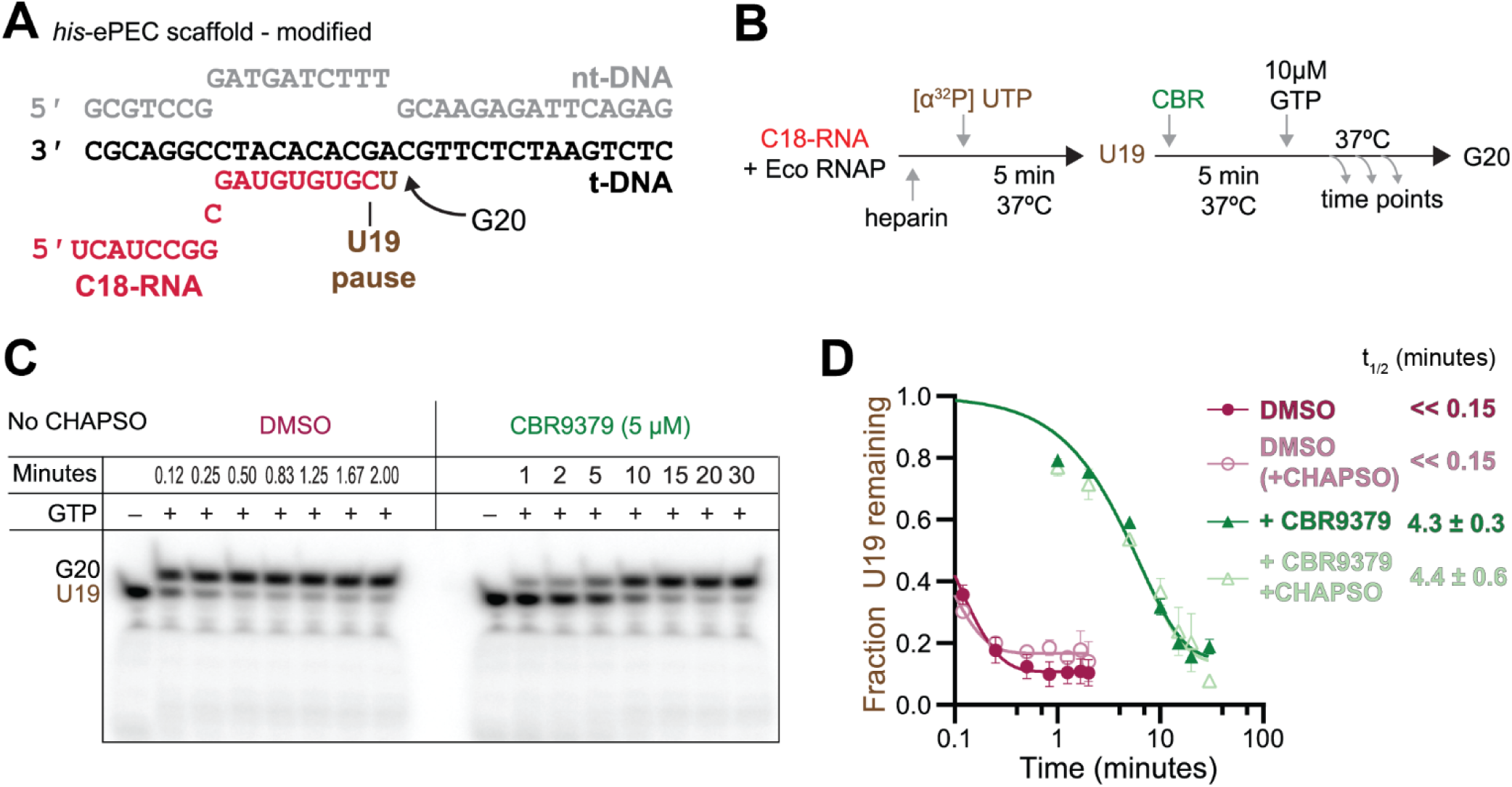
CBR9379 dramatically slows the *Eco*RNAP single-nucleotide addition cycle. (A) The *Eco his*-elemental pause elongation complex (*his*-ePEC) scaffold used for monitoring single nucleotide extension. nt-strand DNA is gray, t-strand DNA is black, RNA is red. (B) Schematic of experimental design to radioactively label C18-RNA in *his*-ePEC scaffold (generating U19), then monitor extension of U19 to G20 as a function of time. (C) Denaturing polyacrylamide gel showing results of single nucleotide extension assay without (DMSO) or with CBR9379. (D) Plot showing the fraction U19 remaining as a function of time without (DMSO -/+ CHAPSO) or with CBR9379 (-/+ CHAPSO). The half-life (t_1/2_) for each condition is shown. Curve fits and half-lives determined using one-phase decay equation (y_0_=1) with GraphPad Prism. Gel image for +CHAPSO is shown in Fig. S2.

In the absence of CBR9379, extension of U19 to G20 was essentially complete after about 1 minute (the t_½_ for 50% conversion of U19 was << 9 seconds; Fig. 2C-D). By contrast, saturating amounts of CBR9379 (5 µM) increased the t_½_ to 4.3 ± 0.3 minutes (258 ± 18 sec; Fig. 2C-D). These findings align with prior observations of reduced transcription speed of *Eco*RNAP in the presence of CBRs (26, 27, 37). The addition of the detergent CHAPSO, used in our cryo-EM sample preparation to mitigate particle orientation bias (38), had no effect on assay kinetics (+CBR9379+CHAPSO t_1/2_ = 4.4 ± 0.6 minutes) (Fig. 2D; Fig. S2E).

### *Eco*RNAP (without CBR) forms an ensemble of *his*-ePEC pause states

We assembled *Eco his*-ePECs and split the sample into two conditions: one with DMSO (–CBR control) and one with a saturating amount of CBR9379. We collected cryo-EM data on each individually and processed each dataset using a similar pipeline (Fig. S3A and S5A; Table S1). In the absence of CBR, we observed three conformations of the active site, which we name Open *Eco*-ePEC (openRH-FL/openTL, 63% of particles, 2.9 Å nominal resolution), Closed *Eco*-ePEC (closedRH-FL/closedTL, 34%, 3.0 Å nominal resolution) and SemiClosed *Eco*-ePEC (openRH-FL/closedTL, 3%, 4.1 Å nominal resolution) (Fig. 3A-C; Fig. S3B, S3C, and S4A-C; Table S1). As observed previously (13), the Open *Eco*-ePEC could be subclassified further into states with different swivel module states (Open^1^, Open^2^, and Open^3^; Fig. S4A-C). In all three Open *Eco*-ePEC classes, the RH-FL domain is rotated out (open), the TL is open and mostly disordered, and SI3 is in an open position (Fig. 3A). By contrast, in Closed *Eco*-ePEC, the RH-FL rotates in toward the active site (closed), the TL is closed, and SI3, as a consequence of TL closing, is in a closed position (Fig. 3B). In the less populated SemiClosed *Eco*-ePEC, the TL and SI3 are closed, but the RH-FL remains open (Fig. 3C).

**Figure 3.**
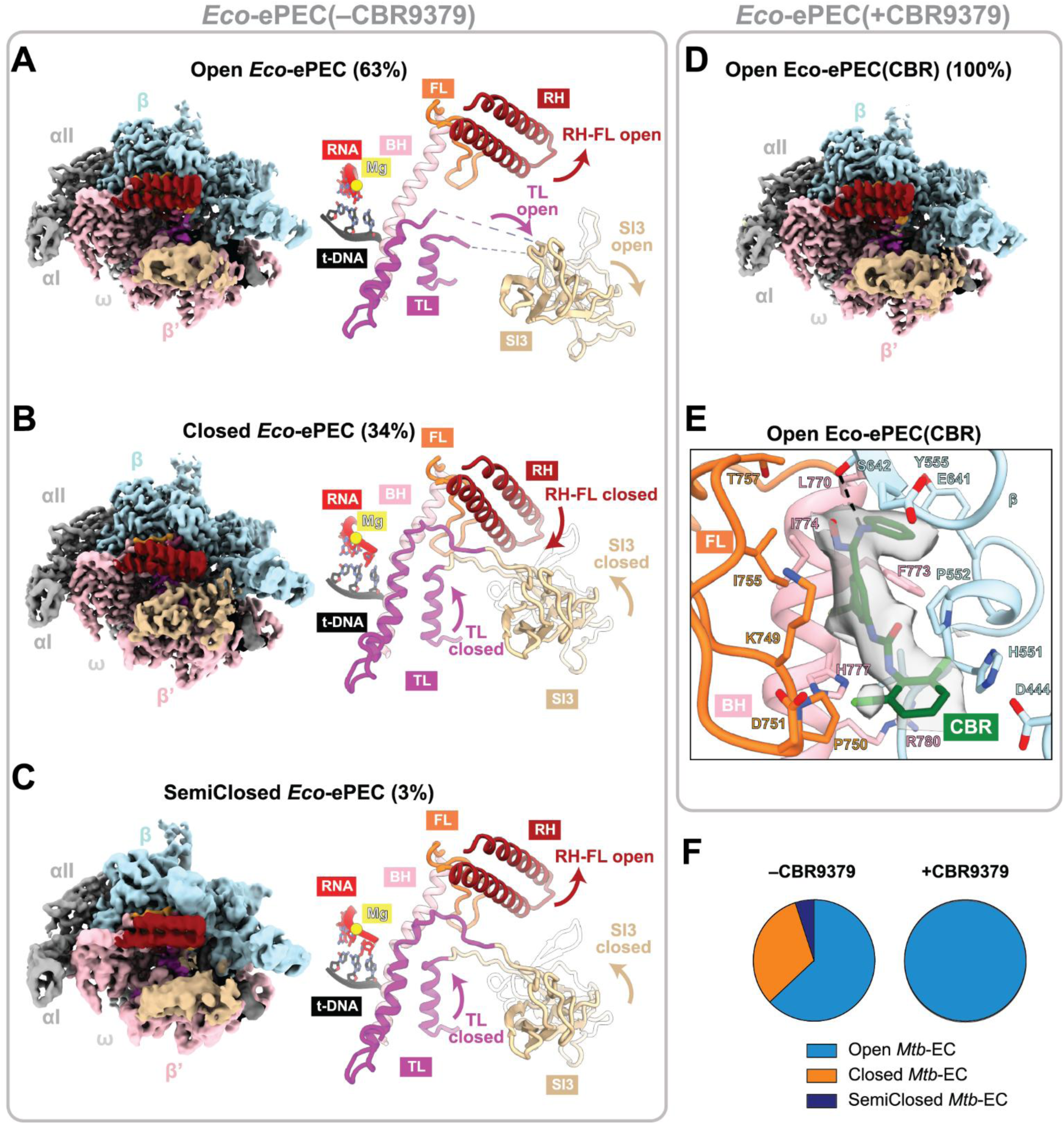
CBR alters the *Eco*-ePECs conformational ensemble and immobilizes the RH-FL in an open state. (A-C) Overall cryo-EM structures and active-site conformations of *Eco*-ePECs. Left: The cryo-EM maps are colored by RNAP subunit or key structural motifs. α and ꞷ are light gray, β light blue, β’ light pink, RH is crimson, FL orange, TL magenta, and SI3 is tan. Right: Close-up and rotated view of active site regions shown using secondary structure cartoons. Nucleic acid color-coding: RNA red, t-DNA black. The RNAP active-site Mg^2+^ is a yellow sphere. Arrows indicate open or closed RH-FL, TL and SI3. (A) Open active-site conformation (RH-FL open, TL open, SI3 open). (B) Closed active-site conformation (RH-FL closed, TL closed, SI3 closed). (C) SemiClosed active-site conformation (RH-FL open, TL closed, SI3 closed). (D) Overall cryo-EM structure Open *Eco*-ePEC(CBR9379). (E) Close-up and rotated view of CBR9379-bound region of *Eco*RNAP. Side chains of residues that interact with CBR9379 (green) are shown. Map density for CBR9379 is shown. (F) Pie charts illustrating cryo-EM particle distributions for *Eco*-ePEC into Open, Closed and SemiClosed active-site states -/+ CBR9379.

Open Eco-ePEC and Closed Eco-ePEC states differ in their nucleic-acid translocation states. The Closed Eco-ePEC is pre-translocated, with the 3′ RNA nucleotide and template-strand DNA (t-DNA) base paired in the *i*+1 site, whereas the Open Eco-ePECs are half-translocated (the 3′ RNA nucleotide has translocated to the *i* site, but the corresponding t-DNA base remains in the *i*+1 site and base paired to the RNA; Fig. S4D and S4E) (13, 19). The pre-translocated Closed Eco-ePEC state is similar to the step immediately after catalysis in the NAC [Fig. 1A(IV)]. The SemiClosed state is likely pre-translocated, but this assignment is tentative due to the low resolution of the cryo-EM reconstruction (Fig. S6F). The half-translocated step is characteristic of some elemental pause complexes (13, 19). These results mirror the *his*-ePEC observations of (13), confirming the robustness of the cryo-EM processing approach.

### CBR holds the *Eco*RNAP RH-FL in an open state

In the presence of CBR9379, the ensemble of *Eco*-ePEC structural states on the *his*-ePEC scaffold (Fig. 3A-C) collapses into two Open states, differing only in swivel module states (Open^1^ and Open^2^; Fig. S5; we name this state Open *Eco*-ePEC(CBR)) (Fig. 3D-F; Fig. S5; Table S1). CBR9379 is bound in a hydrophobic pocket formed by the N-terminal end of the BH, the FL and parts of the β subunit with a polar interaction with βS642 (Fig. 3E), as seen in previous crystal structures (27, 28).

### AAP-SO_2_ dramatically slows the *Mtb*RNAP single-nucleotide addition cycle

Although AAPs and CBRs are chemically unrelated and do not cross-inhibit bacteria, they occupy similar binding pockets on the *Mtb* and *Eco*RNAPs, respectively (27, 28, 32, 33) (Fig. 1E). Therefore, we hypothesized that their mechanisms of action could also be similar. We tested the direct effect of AAP-SO_2_ (Fig. 1D) on *Mtb*RNAP single nucleotide addition (Fig. 4) using the *his*-ePEC scaffold (Fig. 2A). In the absence of AAP-SO_2_, extension of U19 to G20 was essentially complete after about 1 minute; the t_½_ for 50% conversion of U19 was << 9 secs (Fig. 4B and 4C). By contrast, saturating amounts of AAP-SO_2_ (10 µM) increased the t_½_ to 8.4 ± 0.7 minutes (504 ± 42 sec; Figs. 4B and 4C). These findings align with prior observations that AAP-SO_2_ reduced the rate of single nucleotide addition of *Mtb*RNAP (33). The addition of the detergent (1H,1H,2H,2H-perfluorooctyl) phosphocholine (FC8F), used in our cryo-EM sample preparation to mitigate particle orientation bias, had no effect on the assay kinetics (+AAP-SO_2_+FC8F t_½_ = 8.9 ± 1 minutes) (Fig. 4C; Fig. S2E).

**Figure 4.**
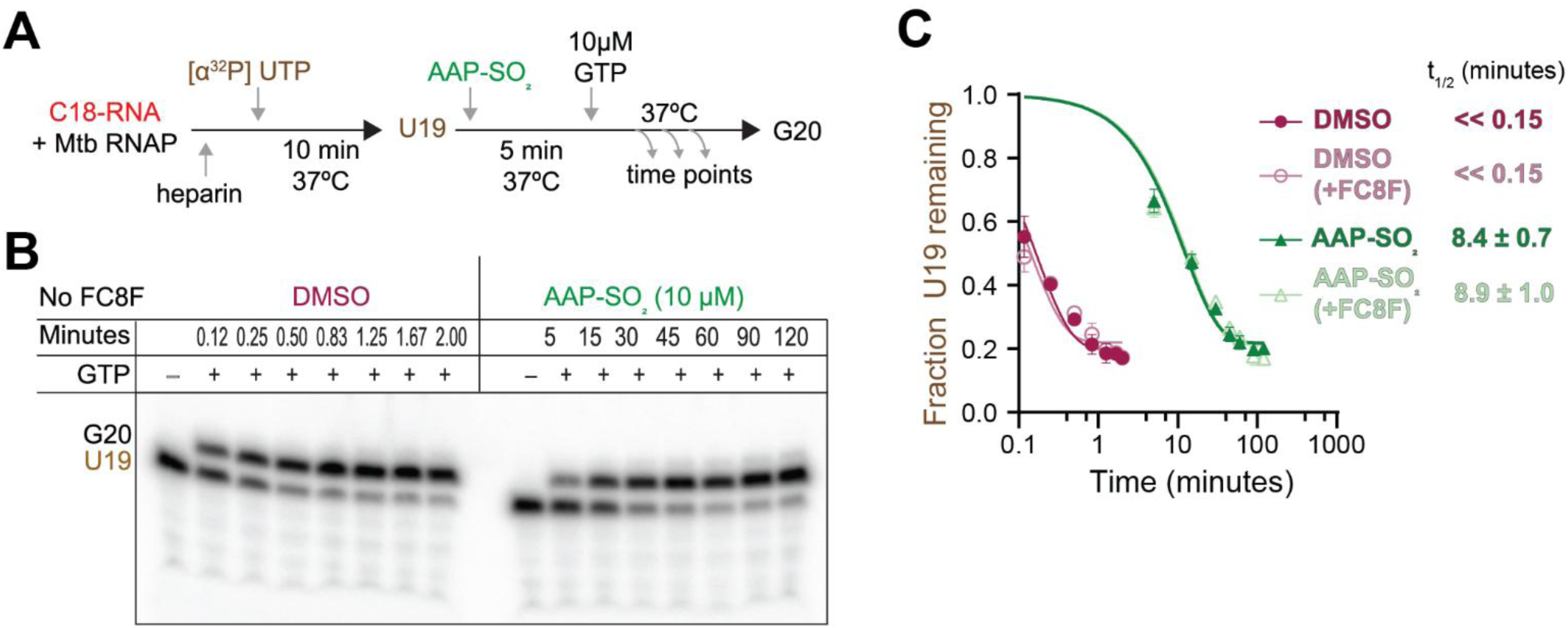
AAP-SO_2_ dramatically slows the *Mtb*RNAP single-nucleotide addition cycle. (A) Schematic of experimental design to radioactively label C18-RNA in *his*-ePEC scaffold (Fig. 2A) to generate U19, then monitor extension of U19 to G20 as a function of time. (B) Denaturing polyacrylamide gel showing results of single nucleotide extension assay without (DMSO) or with AAP-SO_2_. (C) Plot showing the fraction U19 remaining as a function of time without (DMSO -/+ FC8F) or with AAP-SO_2_ (-/+ FC8F). The half-life (t_1/2_) for each condition is shown. Curve fits and half-lives determined using one-phase decay equation (y_0_=1) with GraphPad Prism. Gel image for +FC8F is shown in Fig. S2.

### An *Mtb*RNAP EC forms an ensemble of active-site states

In contrast to *Eco*RNAP, cryo-EM analysis of *Mtb*RNAP on the *his*-ePEC scaffold (Fig. 2A) yielded only an open active-site state (data not shown), consistent with findings that *Mtb*RNAP responds weakly to the *Eco his*-pause signal (39). To obtain an *Mtb*RNAP conformational ensemble that included open and closed active site states, we designed a scaffold to mimic substrate (GTP) loading [Fig. 1A(II)] using an RNA transcript with a 3’-deoxy nucleotide at the 3’-end to prevent catalysis (Fig. 5A; 3’-dG20-RNA).

**Figure 5.**
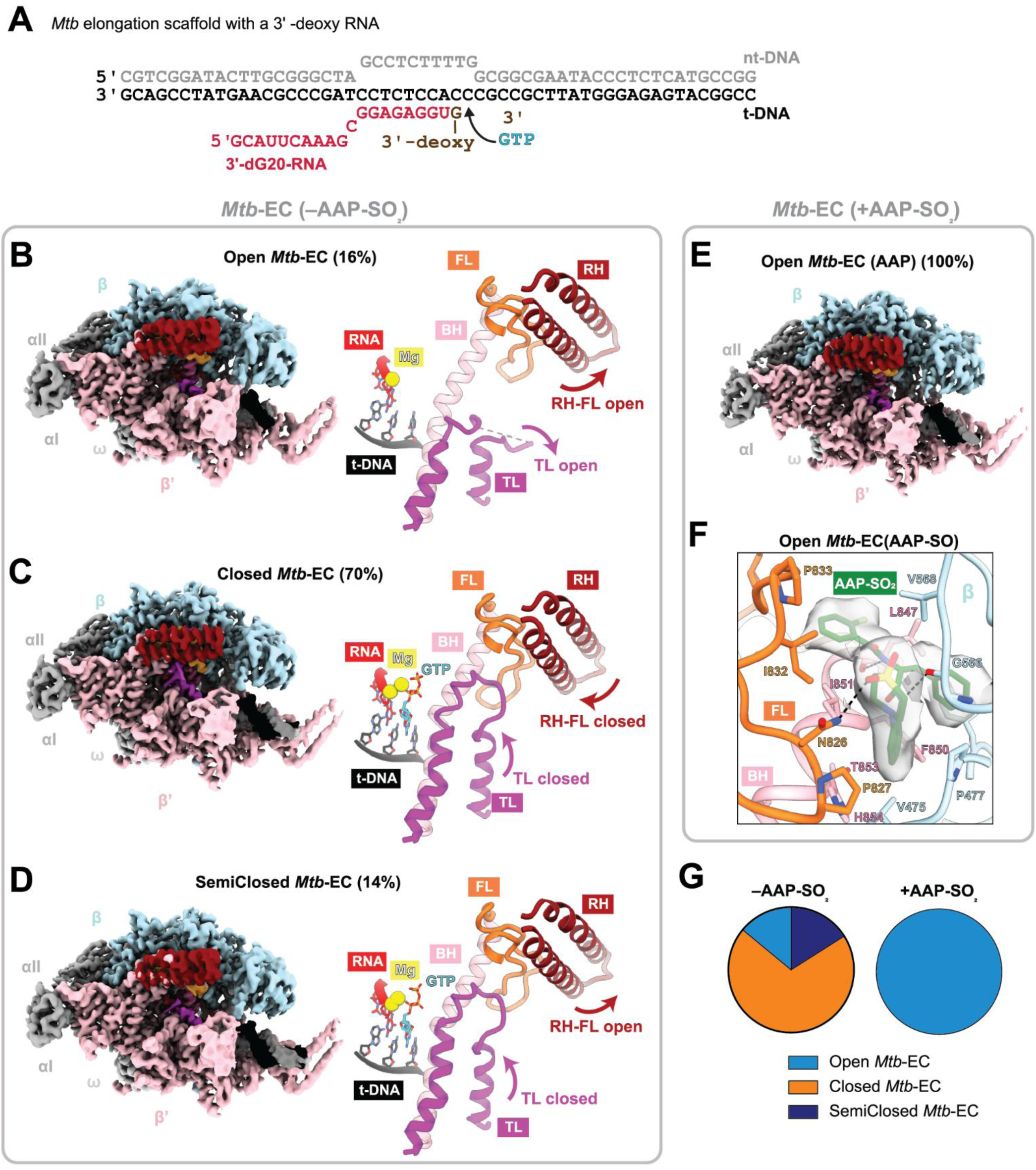
AAP-SO_2_ alters the *Mtb*-ECs conformational ensemble and immobilizes the RH-FL in an open state. (A) Elongation scaffold with 3’-deoxy RNA (3’-dG20-RNA) used for cryo-EM of *Mtb*RNAP. nt-DNA is gray, t-DNA is black, RNA is red. Incoming GTP added during grid preparation is indicated. (B-D) Overall cryo-EM structures and active-site conformations of *Mtb*-ECs. Left: The cryo-EM maps are colored by RNAP subunit or key structural motifs. α and ꞷ are light gray, β light blue, β’ light pink, RH is crimson, FL orange, TL magenta. Right: Close-up and rotated view of active site regions shown using secondary structure cartoons. Nucleic acid color-coding: RNA red, t-DNA black. The RNAP active-site Mg^2+^ is a yellow sphere. Arrows indicate open or closed RH-FL, TL and SI3. (B) Open active-site conformation (RH-FL open, TL open). (C) Closed active-site conformation (RH-FL closed, TL closed). (D) SemiClosed active-site conformation (RH-FL open, TL closed). (E) Overall cryo-EM structure Open *Mtb*-EC(AAP-SO_2_). (F) Close-up and rotated view of AAP-SO_2_-bound region of *Mtb*RNAP. Side chains of residues that interact with AAP-SO_2_ (green) are shown. Map density for AAP-SO_2_ is shown. (G) Pie charts illustrating cryo-EM particle distributions for *Mtb*-EC into Open, Closed and SemiClosed active-site states -/+ AAP-SO_2_.

Cryo-EM with the EC scaffold plus GTP (Fig. 5A) revealed three active site states, Open *Mtb*-EC (16% of the particles, 3.3 Å nominal resolution), Closed *Mtb*-EC (70% of the particles, 3.0 Å nominal resolution), and SemiClosed *Mtb*-EC (14%, 3.3 Å nominal resolution) (Fig. 5B-D; Fig. S7; Table S2). The Closed *Mtb*-EC was characterized by a fully closed TL and closure of the RH-FL domain (Fig. 5C). We observed clear density for the incoming GTP at the i+1 site, consistent with a pre-incorporation intermediate (Fig. S7A). The Open Mtb-EC was in a post-translocated state with an open TL and open RH-FL (Fig. 5D); density for the GTP substrate was absent (Fig. S7B). The GTP substrate was present in the SemiClosed Mtb-EC (Fig. S7C), the TL was closed, but the RH-FL remained open (Fig. 5D).

### AAP-SO_2_ holds the *Mtb*RNAP RH-FL in an open state

In the presence of AAP-SO_2_, the ensemble of *Mtb*-EC structural states on the *Mtb*-EC scaffold (Fig. 5B-D) collapsed into one state, with 100% of the particles resolving into an Open *Mtb*-EC (we name this state Open *Mtb*-EC(AAP-SO_2_); Fig. 5E-G; Fig. S7D and S7E; Table S2). The binding pose and interactions of AAP-SO_2_ match those previously seen in a post-translocated *Mtb*-EC structure bound to AAP-SO_2_ (33) (Fig. 5F).

## DISCUSSION

The CBR and AAP compounds bind ∼30 Å from the RNAP active site Mg^2+^-ion but inhibit all known catalytic activities of the enzyme (26–28, 32, 33). The compounds do not affect NTP substrate binding and do not interfere with translocation (26, 27, 37, 39), indicating an allosteric inhibitory mechanism.

The CBRs have been shown to inhibit *Eco*RNAP TL closing (4, 27), but do so allosterically without contacting the TL (27, 28). We establish here that CBR9379 directly contacts the *Eco*RNAP FL and immobilizes it (Fig. 3). Similarly, AAP-SO_2_ directly contacts the *Mtb*RNAP FL and immobilizes it (Fig. 5). Both compounds thus prevent closure of the RH-FL domain that appears to be coordinated with TL closing in their respective RNAPs [see Supplementary Video S1 (*Eco*RNAP) and Supplementary Video S2 (*Mtb*RNAP)].

Closure of the RH-FL domain onto the closed TL allows the formation of specific FL-TL interfaces (Fig. 6). In the active-site Closed *Eco*-ePEC and *Mtb*-EC structures, van der Waals and hydrogen-bond contacts form between the closed TL and closed RH-FL that are lost in the SemiClosed structures (where the TL is closed but the RH-FL is open; Fig. 6A-D; Fig. S1; Table 1).

**Figure 6.**
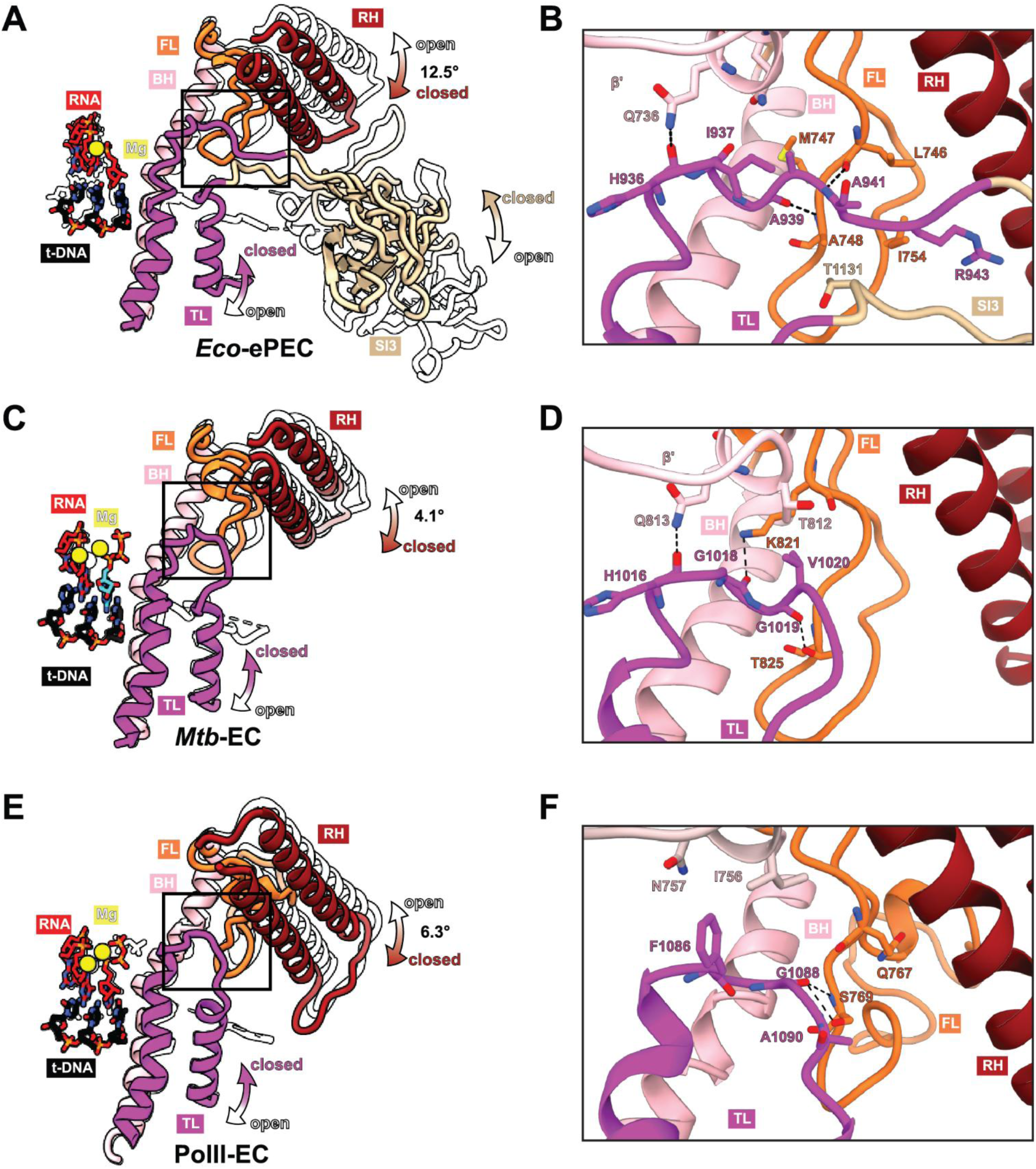
FL-TL interactions of *Eco*, *Mtb*, and *S. cerevisiae* RNAPs. (A, C, E) Overlay of Open and Closed active site regions. Open structures are outlined in black; Closed structures are shown in secondary structure cartoons (RH, crimson; FL, orange; BH, light pink; TL, magenta; SI3 when present, tan; RNA, red; t-DNA, black; active-site Mg^2+^, yellow sphere). Arrows indicate open and closed RH-FL, TL and SI3. Angle for RH-FL rotation is noted. Box indicates region shown in B, D, and F. (B, D, F) Close up, rotated view of interactions between closed RH-FL and closed TL in Closed active site structures. Residues forming H-bonds (gray dashed lines) and Van der Waals interactions are shown. (A, B) *Eco*-ePEC. (C, D) *Mtb*-EC. (E, F) *S. cerevisiae* PolII-EC (active site Open and Closed; PDBs 9RYB and 9QEB, respectively).

**Table 1.** FL-TL interactions in Closed vs. SemiClosed *Eco*-ePEC and Closed vs. SemiClosed *Mtb*-EC.

| *Eco* |  |  |  |
| --- | --- | --- | --- |
| Closed *Eco*-ePEC |  | SemiClosed *Eco*-ePEC |  |
| interacting residues |  |  |  |
| FL | TL | FL | TL |
| van der Waals contacts ($\leq 4.4$ Å) | | | |
| L746 | G939 | - |  |
| M747 | A940 |  |  |
| A748 | A941 |  |  |
| G752 | R943 |  |  |
| I754 | T1131 |  |  |
| Hydrogen bonds ($\leq 3.4$ Å) | | | |
| Q736(NE2) | H936(O) | Q736(NE2) | H936(O) |
| L746(O) | A941(N) | - |  |
| A748(N) | G939(O) | - |  |
| *Mtb* |  |  |  |
| Closed *Mtb*-EC |  | SemiClosed *Mtb*-EC |  |
| interacting residues |  |  |  |
| FL | TL | FL | TL |
| van der Waals contacts ($\leq 4.4$ Å) | | | |
| T812 | G1018 | T812 | V1020 |
| K821 | V1020 | K821 |  |
| Hydrogen bonds ($\leq 3.4$ Å) | | | |
| Q813(NE2) | H1016(O) | - |  |
| K821(NZ) | G1018(O) | - |  |
| T825(OG1) | G1019(O) | T825(OG1) | G1019(O) |

*Eco*RNAP with a TL deletion (DTL-RNAP) catalyzes phosphodiester bond formation about four orders of magnitude more slowly than wild-type RNAP (7, 8). TL mutations expected to inhibit TL folding (inhibit closing) have similar dramatic effects slowing the NAC, while mutations expected to favor TL closing can produce ’fast’ RNAPs, indicating that stimulation of catalysis occurs through the closed TL state (3, 5–8, 20, 40–45). We propose that formation of the specific FL-TL interface upon RH-FL closure stabilizes the closed TL state, increasing the lifetime of the closed TL state and thus allowing more time for the precise configuration of the protein active site and alignment of the incoming NTPs and Mg^2+^-ions for the S_N_2 reaction to proceed (46). Our structural analyses establish that CBR9379/AAP-SO_2_ prevent RH-FL closure, thereby preventing the formation of the FL-TL interface and decreasing the lifetime of the closed TL state. Thus, rapid cycles of TL closing and opening can occur without associated catalysis in the absence of the FL-TL stabilization, slowing the NAC. Single-molecule experiments indicate that an AAP derivative causes prolonged periods of slow *Mtb*RNAP nucleotide incorporation (5 to 50-fold slower than in the absence of AAP) but does not slow the NAC by orders of magnitude (like DTL-RNAP) (47), consistent with the notion that TL closing and the associated stimulation of catalysis can still occur but at a much reduced efficiency (16).

In conclusion, our results establish the structural mechanism for RNAP inhibition by the CBR/AAP compounds. The compounds slow the RNAP NAC by immobilizing the FL, thereby preventing closure of the RH-FL domain that appears to be coordinated with TL closing. The closed FL establishes contacts with the closed TL that stabilize the closed TL state (Fig. 6A-D; Table 1), promoting catalysis. By immobilizing the open FL, the FL-TL contacts cannot form (Table 1) and the closed TL state is relatively destabilized. From these results we infer that RH-FL closure is a crucial RNAP conformational change for efficient progression through the NAC. Because this mechanism is observed in two evolutionarily divergent organisms (*Eco* and *Mtb*) with divergent FL sequences (Fig. S1), and because the TL and RH-FL are conserved structural modules present in all cellular RNAPs across the three domains of life (Fig. S2A-D), we propose that it represents a fundamental and universal mechanism of multi-subunit RNAP function (Fig. 6; Fig. S2A-D).

Establishing the importance of RH-FL motions for efficient RNAP catalysis provides insights for mechanistic design of CBR- and AAP-type inhibitors that inhibit the RH-FL open-to-closed transition.

Moreover, every round of the NAC involves the RH-FL open-to-closed transition, but also the closed-to-open transition to facilitate TL opening, which is required for post-catalysis PPi release and further progression through the NAC (Fig. 1A). Thus, this work defines the closed RH-FL as a new target for antibiotic design. Indeed, any small molecule that interferes with RH-FL motions, in either direction, is predicted to inhibit the RNAP.

## METHODS

### *Eco* RNAP constructs, expression and purification

Full-length *Eco* RNAP comprising rpoA (α), rpoB (β), rpoC (β’), and rpoZ (ω) was expressed and purified following the procedure described previously (48). A pET-based plasmid expressing full-length RNAP subunits α, β, ω and β’-PPX-His10 (PPX; PreScission protease site, LEVLFQGP, Cytiva) under an IPTG-inducible promoter (pVS11 – addgene #128940) was co-transformed with pACYC-Duet-1 encoding ω (addgene #128837) into *Eco* BL21(DE3) and plated on LB agar supplemented with 100 μg/ml ampicillin and 34 μg/ml chloramphenicol, a single colony was used to start an overnight culture at 37°C and 150 RPM. 8 L of LB media supplemented with 100 μg/ml ampicillin and 34 μg/ml chloramphenicol were inoculated with overnight culture (10 mL overnight culture/L of media). The cultures were grown at 37°C to an OD 600 nm of 0.5. The temperature was reduced to 30°C and expression was induced with isopropyl b-D-1-thiogalactopryaoside (IPTG) at a final concentration of 1 mM, and cultures were incubated for an additional 3 h at 30 °C. Cells were harvested by centrifugation at 4000 RCF at 4 °C for 30 min (Beckman JS-4.2). Pellets were resuspended in a total of 120 mL lysis buffer [50 mM Tris-HCl, pH 8.0, 10 mM DTT, 1 mM ZnCl_2_, 5% (v/v) glycerol, 0.5x c0mplete EDTA-free protease inhibitors (Roche), 1 mM PMSF]. Resuspended cells were frozen in liquid N_2_ and stored at −80°C. Cell suspensions were thawed on ice. 0.5x c0mplete EDTA-free protease inhibitors (Roche) and 0.2 mM PMSF were added upon thawing. Cells were lysed using a French press by high pressure shearing (Avestin Emulsi Flex C50). Lysate was centrifuged at 4000 RCF at 4 °C for 30 min (Beckman JA-20) to remove insoluble material. Supernatant from centrifugation was transferred to a beaker and stirred at 4 °C. While stirring, polyethyleneimine [PEI; 10% (w/v) in water adjusted to pH 8.0 with HCl; Fisher Scientific AC178572500] was slowly added to a final concentration of 0.6% (w/v), then the mixture was stirred for 20 min at 4 °C. The PEI precipitate was then collected by centrifugation at 30,000 RCF at 4 °C for 45 min (Beckman JA-20). The pellets were washed three times using 60 mL PEI wash buffer for each wash [50 mM Tris-HCl, pH 8.0, 0.5 M NaCl, 5% (v/v) glycerol, 10 mM DTT]. For each wash, the pellets were resuspended using two 40 ml glass Dounce homogenizers (Wheaton) and the PEI precipitate was collected by centrifugation at 30,000 RCF at 4 °C for 15 min (Beckman JA-20), saving the supernatant for analysis by SDS-PAGE. To elute RNAP from the PEI, the pellets from the last wash step were resuspended as above in 40 ml of PEI elution buffer [50 mM Tris-HCl, pH 8.0, 1 M NaCl, 5% (v/v) glycerol, 10 mM DTT] and the PEI precipitate was collected again by centrifugation. Elution was repeated for two more rounds. After analysis by SDS-PAGE, supernatants from the first two rounds of PEI elution were pooled and transferred to a beaker and stirred slowly at 4 °C. Ammonium sulfate [(NH_4_)_2_SO_4_] was ground to a fine powder using a coffee grinder, then added to the pooled PEI elution slowly while stirring to a final concentration of 35 g/100 mL initial PEI eluate, and the mixture was incubated overnight at 4 °C. Ammonium sulfate precipitate was recovered by centrifugation at 30,000 RCF at 4 °C for 45 min (Beckman JA-18). Pellets were resuspended in 40 mL IMAC buffer A [20 mM Tris-HCl, pH 8.0, 1 M NaCl, 5% (v/v) glycerol, 1 mM β-mercaptoethanol] and nutated at 4 °C for 20 minutes. HiTrap IMAC HP columns (2 x 5 mL; Cytiva) charged with Ni^2+^ were connected in series and equilibrated with IMAC buffer A. Resuspended pellets in IMAC buffer A were filtered through a 5 µm syringe filter and the filtrate was applied on the equilibrated IMAC columns. The columns were then washed with IMAC buffer A containing 0, 25 and 80 mM imidazole consecutively. Protein was eluted in IMAC buffer B [20 mM Tris-HCl, pH 8.0, 1 M NaCl, 250 mM imidazole, 5% (v/v) glycerol, 1 mM β-mercaptoethanol]. Fractions were evaluated by SDS-PAGE and measured for absorbance at 280 nm and pooled according to protein content and purity. His-tagged PPX protease was added at 1:20 molar ratio. The sample was dialyzed overnight at 4 °C in a 12–14 kDa molecular weight cut-off membrane (SpectraPor) against 20 mM Tris-HCl, pH 8.0, 1 M NaCl, 5% (v/v) glycerol, 0.1 mM EDTA, 1 mM β-mercaptoethanol, 0.5 mM DTT. After dialysis, the sample was passed over the IMAC columns followed by dialysis of the flow-through for 4 hours at 4 °C in a 12–14 kDa molecular weight cut-off membrane (SpectraPor) against Biorex A buffer [10 mM Tris-HCl, pH 8.0, 100 mM NaCl, 0.1 mM EDTA, 5% (v/v) glycerol, 5 mM DTT]. After dialysis, the sample was loaded on a 40 mL Biorex column (Biorad Biorex-70 resin #142-5842) equilibrated in Biorex A buffer and eluted with a gradient of 20–80% Biorex B buffer [10 mM Tris-HCl, pH 8.0, 1 M NaCl, 0.1 mM EDTA, 5% (v/v) glycerol, 5 mM DTT] over 15 column volumes. Fractions were pooled according to protein content. The pooled fractions were concentrated using a 100 kDa centrifugal filter (Amicon Ultra, EMD Millipore) at 3500 RCF at 4 °C and loaded on a Superdex 200 26/600 320 mL column (Cytiva) equilibrated in gel filtration buffer [10 mM Tris-HCl, pH 8.0, 0.5 M NaCl, 0.1 mM EDTA, 5% (v/v) glycerol, 5 mM DTT]. Peak fractions were pooled, concentrated using 100 kDa centrifugal filters (Amicon Ultra, EMD Millipore) at 3500 RCF at 4 °C and mixed with gel filtration buffer containing 70% (v/v) glycerol to achieve a final concentration of 20% (v/v) glycerol. Aliquots were flash frozen in liquid N_2_ and stored at −80 °C.

### *Mtb* RNAP constructs, expression and purification

Full-length *Mtb* RNAP comprising rpoA (α), rpoB (β), rpoC (β’), and rpoZ (ω) was expressed and purified following the previously described procedure (49). Expression plasmid pMP61, containing a T7 promoter followed by *rpoA*, *rpoZ*, a fused *rpoBC* with a C-terminal His8 tag, was used. pMP61 was transformed into *Eco* Rosetta 2 cells and plated on LB agar supplemented with 50 μg/ml kanamycin and 34 μg/ml chloramphenicol. A single colony was used to start an overnight culture at 37°C and 150 RPM. 8 L of LB media supplemented with 50 μg/ml kanamycin and 34 μg/ml chloramphenicol were inoculated with overnight culture (10 mL overnight culture/L of media). The cultures were grown at 37°C and 150 rpm to an OD 600 nm of 0.3. The temperature was reduced to 25°C and the culture was grown to an OD 600 nm 0.6. Expression was induced with IPTG at a final concentration of 0.1 mM and cultures were incubated overnight at 25 °C. Cells were harvested by centrifugation at 4000 RCF at 4 °C for 30 min (Beckman JS-4.2). Pellets were resuspended in a total of 120 mL lysis buffer [50 mM Tris-HCl, pH 8.0, 1 mM EDTA, 5 mM DTT, 1 mM ZnCl_2_, 5% (v/v) glycerol, 0.5x c0mplete EDTA-free protease inhibitors (Roche), 1 mM PMSF]. Resuspended cells were frozen in liquid N_2_ and stored at −80°C. Cell suspensions were thawed on ice. 0.5x c0mplete EDTA-free protease inhibitors (Roche) and 0.2 mM PMSF were added upon thawing. Cells were lysed using a French press by high pressure shearing (Avestin Emulsi Flex C50). Lysate was centrifuged at 4000 RCF at 4 °C for 30min (Beckman JA-20) to remove insoluble material. Supernatant from centrifugation was transferred to a beaker and stirred at 4 °C. While stirring, polyethyleneimine [PEI; 10% (w/v) in water adjusted to pH 8.0 with HCl] was slowly added to a final concentration of 0.6% (w/v), and the mixture was stirred for 20 min at 4 °C. PEI precipitate was then collected by centrifugation at 30,000 RCF at 4 °C for 45 min (Beckman JA-20). The pellets were washed three times using 60 mL PEI wash buffer for each wash [50 mM Tris-HCl, pH 8.0, 0.5 M NaCl, 5% (v/v) glycerol, 10 mM DTT]. For each wash, the pellets were resuspended using two 40 ml glass Dounce homogenizers (Wheaton) and the PEI precipitate was collected by centrifugation at 30,000 RCF at 4 °C for 15 min (Beckman JA-20), saving the supernatant for analysis by SDS-PAGE. To elute RNAP from the PEI, the pellets from the last wash step were resuspended as above in 40 ml of PEI elution buffer [50 mM Tris-HCl, pH 8.0, 1 M NaCl, 5% (v/v) glycerol, 10 mM DTT] and the PEI precipitate was collected again by centrifugation. Elution was repeated for two more rounds. After analysis by SDS-PAGE, supernatants from the first two rounds of PEI elution were pooled and transferred to a beaker and stirred slowly at 4 °C. Ammonium sulfate [(NH_4_)_2_SO_4_] was ground to a fine powder using a coffee grinder, then added to the pooled PEI elution slowly while stirring to a final concentration of 35 g/100 mL initial PEI eluate, and the mixture was incubated overnight at 4 °C. Ammonium sulfate precipitate was recovered by centrifugation at 30,000 RCF at 4 °C for 45 min (Beckman JA-18). Pellets were resuspended in 40 mL IMAC buffer A [20 mM Tris-HCl, pH 8.0, 1 M NaCl, 5% (v/v) glycerol, 1 mM β-mercaptoethanol] and nutated at 4 °C for 20 minutes.

HiTrap IMAC HP columns (2 x 5 mL; Cytiva) charged with Ni^2+^ were connected in series and equilibrated with IMAC buffer A. Resuspended pellets in IMAC buffer A were filtered through a 5 µm syringe filter and the filtrate was applied on the equilibrated IMAC columns. The columns were then washed with IMAC buffer A containing 0, then 50 mM imidazole consecutively. Protein was eluted in IMAC buffer B [20 mM Tris-HCl, pH 8.0, 1 M NaCl, 250 mM imidazole, 5% (v/v) glycerol, 1 mM β-mercaptoethanol]. Fractions were run on SDS-PAGE and measured for absorbance at 280 nm and pooled according to protein content and purity. The pooled fractions were loaded on a Superdex 200 26/600 320 mL column (Cytiva) equilibrated in gel filtration buffer [10 mM Tris-HCl, pH 8.0, 0.5 M NaCl, 0.1 mM EDTA, 5% (v/v) glycerol, 5 mM DTT]. Peak fractions were pooled and mixed with buffer containing 70% (v/v) glycerol to achieve a final concentration of 20% (v/v) glycerol. Aliquots were frozen in liquid N_2_ and stored at −80 °C.

### Oligonucleotides used for biochemical and structural work

PAGE purified synthetic DNA oligonucleotides were obtained from Integrated DNA Technologies. HPLC purified RNA oligonucleotides were obtained from Horizon Discovery / Dharmacon. Oligonucleotides were dissolved in RNase-free water (Ambion) to 100 – 200 µM concentrations.

### Nucleotide addition assays

In initial experiments with the original *his*-ePEC scaffold used previously (13), we observed backtracking of RNAP, evident from the presence of short nucleolytic cleavage products. This complicated quantification, since cleavage activity is also influenced by CBR concentration (27). We designed a modified *his*-ePEC scaffold to prevent backtracking (allowing for the upstream end of the transcription bubble to reanneal as the RNA extends to G20), leading to the absence of cleavage products and simplified quantification.

Transcription buffer compositions for *Eco* RNAP and *Mtb* RNAP are listed below:

*Eco* RNAP transcription buffer: 25 mM HEPES-KOH, pH 8.0; 130 mM KCl; 5 mM MgCl₂; 0.15 mM EDTA; 5% glycerol (v/v); 25 µg/mL acetylated BSA; 5 mM DTT.

*Mtb* RNAP transcription buffer: 10 mM HEPES-KOH, pH 8.0; 50 mM potassium glutamate, 10 mM magensium acetate, 1 mM EDTA; 5 µg/mL acetylated BSA; 5 mM DTT.

Template DNA strand (t-DNA) and RNA (C18) for modified-*hi*s-ePEC scaffold (Fig. 2A) were annealed at a molar ratio of 1:1.5 in 1× transcription buffer to prepare t-DNA:RNA hybrid. Annealing was performed by heat denaturation followed by slow cooling: 95 °C for 2 min, 75 °C for 2 min, 45 °C for 5 min, 40 °C for 2 min, 35 °C for 2 min, 30 °C for 2 min, and 25 °C for 2 min, with a cooling rate of 0.1 °C/s between each step. The annealed scaffold was stored at −80 °C until further use.

To assemble the *his*-ePEC complex, the t-DNA:RNA hybrid was mixed with RNAP at a molar ratio of 1:1.5:1.5 (t-DNA:RNA:RNAP) and incubated for 15 min at room temperature (RT). Non-template DNA strand (nt-DNA) was then added at a 2× molar excess relative to t-DNA (final molar ratio tDNA:RNA:RNAP:ntDNA = 1:1.5:1.5:2) and incubated for an additional 15 min at RT. Heparin (Sigma-Aldrich H3149) was added to a final concentration of 0.1 mg/mL to trap free RNAP. The assembled *his*-ePEC was adjusted to 400 nM RNAP with transcription buffer and incubated at 37 °C for 5 min. The RNA was isotope-labeled by adding [α-³²P]UTP (33 Ci/mmol) to extend the C18 RNA to U19 (Fig. 2B). Reactions were incubated at 37 °C for 5 min for *Eco* RNAP and 10 min for *Mtb* RNAP. The reactions were then split into two equal portions: one for control/DMSO and the other for inhibitor treatment.

For the *Eco* RNAP reaction, CBR9379 (dissolved in 100% anhydrous DMSO; Fisher Scientific NC0304463) was added to a final concentration of 5 µM (final DMSO concentration 1.6%). For the *Mtb* RNAP reaction, AAP-SO₂ (dissolved in 100% anhydrous DMSO) was added to a final concentration of 10 µM (final DMSO concentration 1.6%). In control reactions, 100% DMSO was added to match the final DMSO concentration of 1.6%. All reactions were incubated at 37 °C for 5 min. An equal volume of 20 µM GTP in transcription buffer was added to each reaction (final concentrations: 10 µM GTP, 200 nM RNAP), followed by incubation at 37 °C. 5 µL of the reaction was transferred to a new tube with 2X stop buffer (8 M urea, 50 mM EDTA, 0.02% bromophenol blue, 0.02% xylene cyanol in 1X TBE buffer) at desired time points. Time points collected are indicated in the gel images and graphs shown in Fig. 2, 4 and Fig. S2E. Samples were heated at 95°C for 5 min and run on denaturing urea-PAGE (15%; 19:1 acrylamide: bis-acrylamide). The gels were exposed for 2-3 h using phosphor screens and imaged on a Typhoon (Cytiva Amersham).

### Quantitation of nucleotide addition assays

ImageJ (50) was used to generate density profiles for each gel lane, and the corresponding x- and y-values for different time points were exported to Microsoft Excel. These values were then imported into IgorPro (WaveMetrics), where the Multipeak Fitting package was used to fit each profile individually using a Gaussian mixture model. This approach effectively separated the intensity signals from U19 and G20 RNA bands, enabling non–user-biased quantitation. Areas under the fitted peaks for U19 and G20 were used to calculate the fraction of U19 remaining in each lane. Values from three replicates were used to generate graphs and fit to a one-phase decay equation in GraphPad Prism (Fig. 2D and 4C).

### Sample preparation for cryo-EM

*<u>Eco his</u>*<u>-ePECs</u>. t-DNA and RNA (U19) for *his*-ePEC scaffold were annealed at a molar ratio of 1:1.5 in buffer (20 mM Tris-HCL, pH 8.0, 150 mM potassium glutamate) to prepare t-DNA:RNA hybrid.

Annealing was performed by heat denaturation followed by slow cooling: 95 °C for 2 min, 75 °C for 2 min, 45 °C for 5 min, 40 °C for 2 min, 35 °C for 2 min, 30 °C for 2 min, and 25 °C for 2 min, with a cooling rate of 0.1 °C/s between each step. The annealed scaffold was stored at −80 °C until further use.

Purified *Eco* RNAP was buffer exchanged using a Superose 6 Increase (Cytiva) column into cryo-EM buffer (20 mM Tris-HCL, pH 8.0, 150 mM potassium glutamate, 5 mM MgCl_2_, 5 mM DTT). *his*-ePEC t-DNA:RNA was added in 1.5x molar excess to the eluted RNAP and incubated at RT for 15 min. nt-DNA was added in 2x molar ratio to t-DNA (final molar ratio tDNA:RNA:RNAP:ntDNA = 1.5:2.25:1:3). Additional MgCl_2_ was added in accordance with the volume of t-DNA:RNA hybrid and nt-DNA was added for a final 5 mM MgCl_2_ concentration. The mix was incubated at RT for 15 min.

Assembled *his*-ePECs were then concentrated using 0.5 mL 100 kDa centrifugal filter (Amicon Ultra, EMD Millipore) at 10,000 RCF at 4 °C. to ∼ 5 mg/mL RNAP before grid preparation. The samples were incubated in ice and equilibrated back to RT for 10 min prior to grid freezing.

*<u>Mtb</u>* <u>ECs</u>. t-DNA and RNA (U19) for elongation scaffold with a 3′ deoxy-RNA (Fig. 5A) were annealed at a molar ratio of 1:2 in annealing buffer (100 mM HEPES, pH 8.0, 0.5 M NaCl, 10 mM EDTA) to prepare t-DNA:RNA hybrid. Annealing was performed by heat denaturation 95 °C for 5 min followed by slow cooling at RT. The annealed scaffold was stored at −80 °C until further use.

*Mtb* RNAP was dialyzed overnight into *Mtb* cryo-EM buffer (20 mM HEPES, pH 7.5, 150 mM potassium glutamate, 5 mM magnesium acetate, 2.5 mM DTT). t-DNA:RNA hybrid was added to the RNAP at a 1.1x molar excess to RNAP and incubated at RT for 15 min. nt-DNA was added in 1.5x molar excess to t-DNA and incubated at RT for 15 min. Final molar ratio t-DNA:RNA:RNAP:nt-DNA was 1:2:0.9:1.5. The sample was concentrated to ≤ 500 µL using a 0.5 mL 3 kDa centrifugal filter (Amicon Ultra, EMD Millipore) at 10,000 RCF at 4 °C. Concentrated sample was loaded on a Superose 6 Increase column (Cytiva) equilibrated with *Mtb* cryo-EM buffer. Peak fractions with fully assembled ECs were pooled and concentrated using 0.5 mL 3 kDa centrifugal filter (Amicon Ultra, EMD Millipore) at 10,000 RCF at 4 °C to ∼ 6 mg/mL of RNAP before grid preparation. The samples were incubated in ice and equilibrated back to RT for 10 min prior to grid freezing.

### Cryo-EM grid preparation

ECs were split into two tubes. To one tube, 25x molar excess of the inhibitor (dissolved in 100% DMSO) was added to a final DMSO percentage of 1.5%. To the other tube, 100% DMSO was added to a final concentration of 1.5 %. The samples were incubated for 5 minutes at RT before freezing.

C-flat holey carbon grids (CF-1.2/1.3-4Au, EMS) were glow-discharged for 5 s at 25 mA and 0.3 mbar air atmosphere (Pelco EasiGlow) before use. Sample application and vitrification was performed using a Vitrobot Mark IV (ThermoFisher Scientific) equilibrated to 22°C and 100% relative humidity in the blotting chamber.

Before freezing, CHAPSO was added to a final concentration of 8 mM for *Eco his*-ePECs (38). Similarly, (1H,1H,2H,2H-perfluorooctyl) phosphocholine (fluorinated fos-choline-8, FC8F; Anatrace #F300F) was added to 1.5 mM final concentration for *Mtb* ECs. Specifically, 3.6 µL sample was mixed with 0.4 µL of 10X detergent (80 mM CHAPSO or 15 mM FC8F) in a separate tube. 3.6 µL of detergent mixed sample was applied per grid, blotted for 3-4 seconds at blot force -1 or +1 and plunge frozen in liquid ethane.

### Cryo-EM data acquisition

*<u>Eco his</u>*<u>-ePECs</u>. Grids were imaged using a 300 kV Titan Krios (ThermoFisher Scientific) equipped with a K3 camera (Gatan), a Cs corrector and a BioQuantum imaging filter (Gatan). Images were recorded using SerialEM with a pixel size of 0.676 Å/px over a nominal defocus range of −0.6 to −2.0 μm and

20 eV energy filter slit width. A total of 26,972 (CBR minus) and 28,709 (CBR plus) gain-normalized movies were recorded in counting mode (K3 camera; image dimensions of 4092 × 5760 px) with 20 e−/px/s (at the camera) in dose-fractionation mode of 0.03 s over a 1.2 s exposure (40 frames) to give a total dose of 52.5 e−/Å^2^.

*<u>Mtb</u>* <u>ECs</u>. Grids were imaged using a 300 kV Titan Krios (ThermoFisher Scientific) equipped with a K3 camera (Gatan). Images were recorded using SerialEM with a pixel size of 0.847 Å/px over a nominal defocus range of −0.6 to −2.0 μm and 20 eV energy filter slit width. A total of 34,503 (AAP-SO_2_ minus) and 28,412 (AAP-SO_2_ plus) gain-normalized movies were recorded in counting mode (image dimensions of 4092 × 5760 px) with 27.5 e−/px/s (at the camera) in dose-fractionation mode of 0.04 s over a 1.4 s exposure (35 frames) to give a total dose of 50.7 e−/Å^2^.

### Cryo-EM data processing

Most structural biology software was accessed through the SBGrid software package (51). All processing was performed using cryoSPARC v4 (52). Movies were drift-corrected and summed using Patch Motion Correction. The contrast transfer function (CTF) was estimated for each micrograph using Patch CTF Estimation. Micrographs with estimated CTF fit resolutions worse than 10 Å were excluded. Particles were picked using the CS4 blob picker (particle diameter: 100–300 Å) and extracted with a box size of 384 px. Extracted particles were Fourier-cropped to 128 px and subjected to two rounds of 2D classification.

<u>No CBR *Eco*-ePECs (Fig. S3A)</u>. 2D classification yielded 2,324,608 particles. Two initial models were generated using ab initio reconstruction from a subset of 1,000,000 particles, resulting in one RNAP-like and one junk class. The full dataset was further curated through two rounds of heterogeneous refinement using the ab initio classes, with the junk class included as a decoy. Particles contributing to RNAP-like reconstructions (1,675,000 particles) were combined, re-extracted with a 384 px box, and subjected to Non-Uniform (NU) refinement (53). Overrepresented orientations were removed using Rebalance Orientations, resulting in 978,427 particles.

Particle subtraction was performed using a mask around the RH-FL domain, followed by 3D classification (n=4). Reconstructed volumes were manually inspected. Particles from one class representing a closed *Eco*-ePEC state were subjected to NU refinement with defocus and global CTF refinement enabled (54), followed by reference-based motion correction (RBMC) (55), additional NU refinement, and local resolution estimation to generate a locally filtered map. Particles from another reconstruction exhibiting anisotropy and low resolution were separated and used for heterogeneous refinement (n=2), producing both open and closed states; these were excluded from further analysis. Particles from the remaining two classes (representing the open state) were pooled, processed through RBMC, and subjected to heterogeneous refinement. After manual inspection, one reconstruction representing a SemiClosed state with closed SI3 was subjected to local resolution estimation and locally filtered map generation. For the remaining particles, particle subtraction was performed to retain only the SI3 domain signal.

The remaining particles all representing an Open state were used in 3D classification, classified into sub-classes due to swiveling (13, 19) (Fig. S3A and S4A-C). The swivel module for *Eco*RNAP includes the clamp, jaw, SI3, dock and jaw and makes about a third of the total RNAP mass. The PDB 8EHI model was rigid body fit using Phenix into each of the volumes. The rotation of the swivel module for each model was determined using the draw_rotation_axis PyMOL script. Particles from 6 classes were pooled to generate three classes, Open^1^ *Eco*-ePEC, Open^2^ *Eco*-ePEC and Open^3^ *Eco*-ePEC from here on, with swivel angles are 2.42°, 3.65° and 4.58°, consistent with previously observed range of angles for *his*-ePECs structures with an open active site. Swiveling is characteristic of paused transcription complexes and is believed to play a role in preventing forward translocation of the polymerase, thereby resulting in a paused state (13, 19). The nominal resolutions for Closed Eco-ePEC, Open^1^ *Eco*-ePEC, Open^2^ *Eco*-ePEC, Open^3^ *Eco*-ePEC and SemiClosed *Eco*-ePEC are 3.0 Å, 2.9 Å, 3.3 Å, 2.7 Å and 4.1 Å (Fig. S3B, S3C, S4A-C).

<u>CBR plus *Eco*-ePECs (Fig. S5A).</u> 2D classification yielded 2,950,656 particles. Two initial models were generated using ab initio reconstruction from a subset of 1,000,000 particles, resulting in one RNAP-like and one junk class. The full dataset was further curated through two rounds of heterogeneous refinement using the ab initio classes, with the junk class included as a decoy. Particles contributing to RNAP-like reconstructions (1,084,014 particles) were combined, re-extracted with a 384 px box, and subjected to Non-Uniform (NU) refinement. Overrepresented orientations were removed using Rebalance Orientations, resulting in 622,144 particles.

Particle subtraction was performed using a mask around the RH-FL domain, followed by 3D classification (n=4). Reconstructed volumes were manually inspected. Particles from three volumes were pooled and used for heterogeneous refinement. Particles classifying into a low resolution volume were removed from further analysis. 318,790 particles were subjected to RBMC and used for 3D variability analysis (56) to examine swiveling. Swivel angles for the dataset ranged from 2.2° to 4.64°. Particles were separated into 4 clusters using 3D variability display, resulting in four volumes, one with 19,250 particles with a 4.64° swivel angle, 146,299 particles with a 3.84° swivel angle, 127,272 particles with a 2.75° swivel angle, and 16,972 particles with a 2.19° swivel angle. Particles from cluster 1 and 2 were pooled to generate Open^2^ *Eco*-ePEC(CBR) with nominal resolution 2.9 Å (Fig. S5B). Particles from cluster 3 and 4 were pooled to generate Open^1^ *Eco*-ePEC(CBR) with nominal resolution 2.9 Å (Fig. S5C).

<u>AAP minus *Mtb*-ECs (Fig. S6A)</u>. 2D classification yielded 733,444 particles. Two initial models were generated using ab initio reconstruction from a subset of 1,000,000 particles, resulting in one RNAP-like and one junk class. The full dataset was further curated through two rounds of heterogeneous refinement using the ab initio classes, with the junk class included as a decoy. Particles contributing to RNAP-like reconstructions (520,654) were combined, re-extracted with a 384 px box, and subjected to Non-Uniform (NU) refinement to generate a consensus map.

Particle subtraction was performed using a mask around the RH-FL domain, followed by 3D classification (n=4). Reconstructed volumes were manually inspected. Particles from two classes representing a closed *Mtb*-EC state were subjected to NU refinement with defocus and global CTF refinement enabled, followed by RBMC, additional NU refinement, and local resolution estimation to generate a locally filtered map.

Particles from another reconstruction resembling a closed *Mtb*-EC exhibiting low resolution were excluded from further analysis. Particles from the remaining class were processed through RBMC and subjected to a round of heterogeneous refinement (n=2), yielding one class with an open *Mtb*-EC and another with a SemiClosed *Mtb*-EC. Each class was NU Refined, used for local resolution determination and local filtering. Three volumes for AAP minus *Mtb*-ECs were generated in total, Closed *Mtb-*EC, SemiClosed *Mtb-*EC and Open Mtb-EC with nominal resolutions 3.0 Å, 3.3 Å and 3.3 Å, respectively (Fig. S6B-D).

<u>AAP plus Mtb-ECs (Fig. S7D)</u>. 2D classification yielded 941,367 particles. The full dataset was further curated through two rounds of heterogeneous refinement using the AAP minus *Mtb*-EC volume and junk volume included as a decoy. Particles contributing to RNAP-like reconstructions (453,888) were combined, re-extracted with a 384 px box, and subjected to Non-Uniform (NU) refinement to generate a consensus map.

Particle subtraction was performed using a mask around the RH-FL domain, followed by 3D classification (n=4). Reconstructed volumes were manually inspected. Particles from two classes representing a open *Mtb*-EC state were subjected to NU refinement with defocus and global CTF refinement enabled, followed by reference-based motion correction (RBMC), additional NU refinement, and local resolution estimation to generate a locally filtered map.

Particles from another reconstruction resembling an Open *Mtb*-EC exhibiting low resolution were excluded from further analysis. Particles from the remaining class did not look like RNAP and were excluded.

### Model building, refinement and validation

Initial models were generated by fitting the structures 8EHI (13) (Open Eco-PEC), 8EH8 (13) (Closed Eco-PEC), and 8E95 (49)(Open Mtb-EC) into the corresponding cryo-EM density maps using ChimeraX (57). Each model was subjected to rigid-body refinement with phenix.real_space_refine (58) against its respective map to obtain initial models for analysis. The refined models were then analyzed and modified in ChimeraX and Coot (59). phenix.real_space_refine was used to finalize all models, including all atom refinement and B-factor refinement, with Ramachandran and secondary-structure restraints, as well as restraints for CBR9379 and AAP-SO₂.

### Data, Materials, and Software Availability

All unique/stable reagents generated in this study are available without restriction from the lead contact, S.A.D. The cryo-EM density maps and atomic coordinates have been deposited in the EMDataBank and PDB as follows: Open^1^ *Eco*-ePEC (EMD-75280, PDB 10MB), Open^2^ *Eco*-ePEC (EMD-75281, PDB 10MC), Open^3^ *Eco*-ePEC (EMD-75282, PDB 10MD), Closed *Eco*-ePEC (EMD-75279, PDB 10MA), SemiClosed *Eco*-ePEC (EMD-75283, PDB 10ME), Open^1^ *Eco*-ePEC(CBR9379) (EMD-75284, PDB 10MF), Open^2^ *Eco*-ePEC(CBR9379) (EMD-75285, PDB 10MG), Open *Mtb*-EC (EMD-75287, PDB 10MJ), Closed *Mtb*-EC (EMD-75286, PDB 10MI), SemiClosed *Mtb*-EC (EMD-75288, PDB 10MK), Open *Mtb*-EC(AAP-SO_2_) (EMD-75289, PDB 10ML). The models used for initial building (PDBs 8EH8, 8EHA, 8EHI, 8E95, 9MRQ) are available in the PDB.

### Author Contributions

Y.D., R.L., E.A.C., and S.A.D. conceived and designed this study. Y.D. performed protein purification and biochemistry, prepared cryo-EM specimens and processed all cryo-EM data. Y.D. and S.A.D. built and analyzed atomic models. R.L., E.A.C., and S.A.D. supervised and acquired financial support. Y.D. and S.A.D. wrote the first draft of the manuscript; all authors contributed to the final version.

## Acknowledgements

We thank Markus Lang and Adrian Richter (Institute of Pharmacy, Martin-Luther-University Halle-Wittenberg, Halle, Germany) for providing AAP-SO_2_, Qingrong Li and Dong Wang (University of California San Diego) for sharing *S. cerevisiae* PolII cryo-EM structures before publication (PDBs 9RYB and 9QEB), and members of the Darst and Campbell laboratories for helpful discussions, especially Andreas Mueller for advice on cryo-EM data processing. We thank Johanna Sotiris, Honkit Ng, and Mark Ebrahim of the Evelyn Gruss Lipper cryo-EM Resource Center (The Rockefeller University) for support with electron microscopy instrumentation and sample handling.

Some of this work was performed at the Simons Electron Microscopy Center at the New York Structural Biology Center, with major support from the Simons Foundation (SF349247). Y.D. was supported by the Pels Family Center for Biochemistry and Structural Biology as part of a grant from the Donald A. Pels Charitable Trust to The Rockefeller University. This research was supported by NIH grants R01 GM38330 to R.L., R35 GM151879 to E.A.C., and R35 GM118130 to S.A.D.

## Supplementary Materials

**Figure S1.**
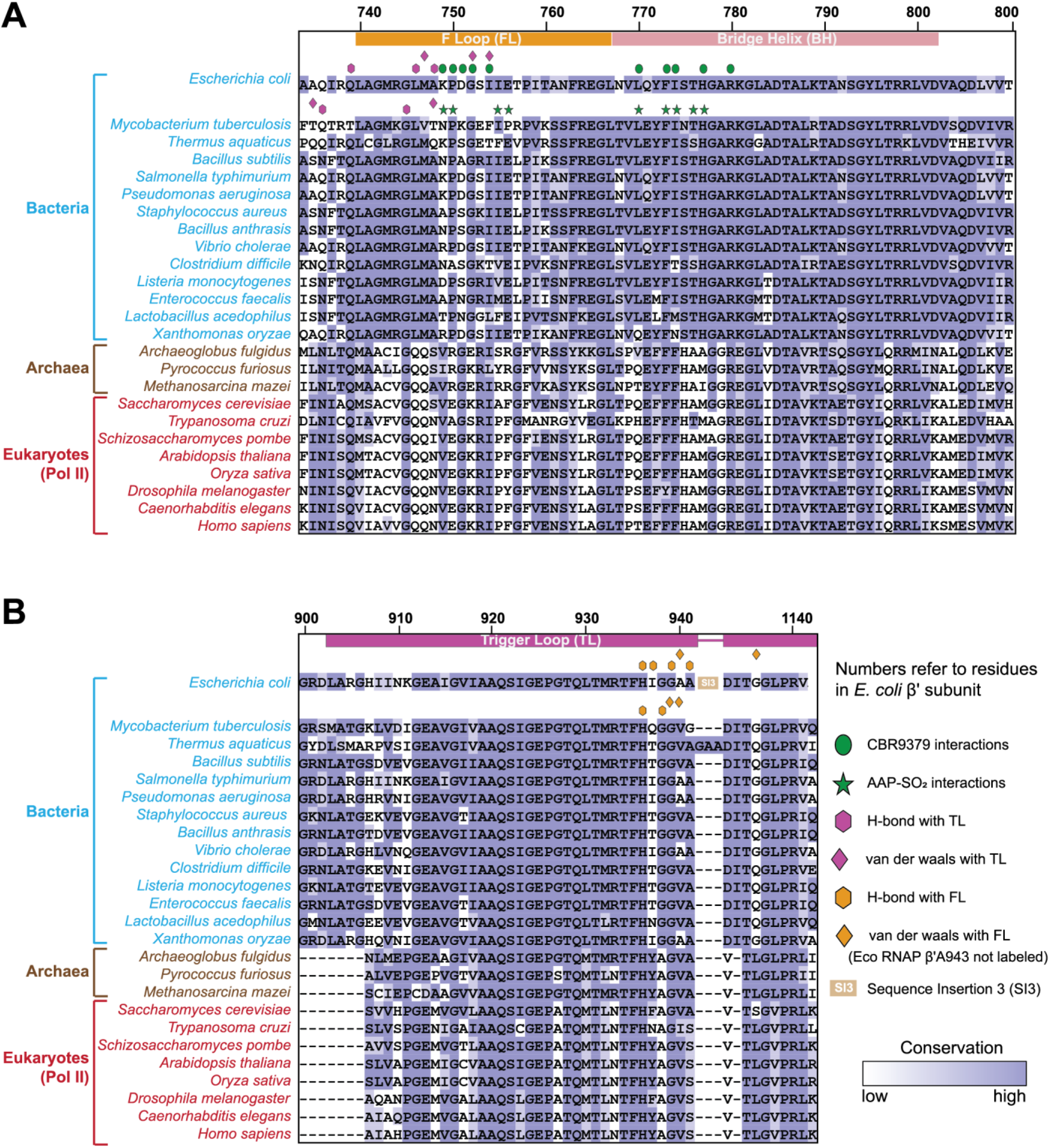
Conservation of BH, FL and TL conservation across domains of life. (A) Sequence alignment of rpoC (β’) or equivalent subunit of RNAP across species for FL (orange bar on top) and BH (light pink bar on top). Bacteria are labeled in blue, Archaea in brown, and eukaryotic PolII in red. Sequence numbering at top is shown for *Eco*RNAP β’. Green ellipses and stars indicate residues that interact with CBR9379 and AAP-SO_2_, respectively. Pink hexagons (H-bonds) and rhombi (Van der Walls) indicate interactions with TL. (B) Sequence alignment of rpoC (β’) or equivalent subunit of RNAP across species for TL residues (magenta bar on top) shown. Bacteria are labeled in blue, Archaea in brown and eukaryotic PolII in red. Sequence numbering at top is shown for *Eco*RNAP β’. Orange hexagons (H-bonds) and rhombi (Van der Waals) indicate interactions with FL residues.

**Figure S2.**
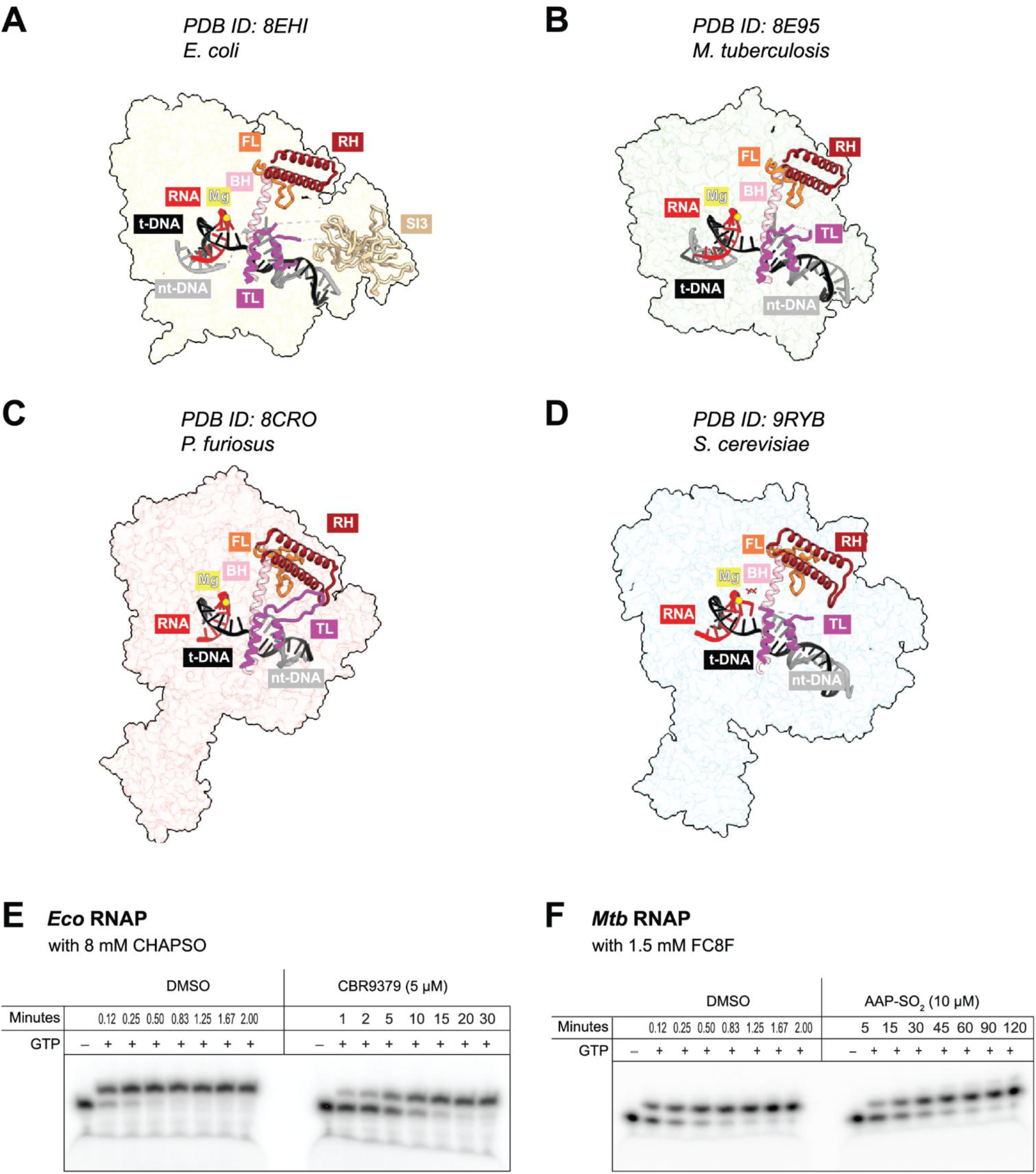
RNAP architecture across species; CBR9379 and AAP-SO_2_ activities in the presence of detergents. (A-D) Surface views of *Eco, Mtb, P. furiosus and S. cerevisiae* RNAP structures, with RH, FL, BH, TL, *Eco*RNAP-SI3, RNA, t-DNA and nt-DNA shown in secondary structure cartoon. Active site Mg^2+^ (yellow) is shown in spheres. (A) Bacteria: *Eco his*-ePEC [PDB 8EHI; (1)].(B) Bacteria: *Mtb*-EC [PDB 8E95; (2)]. (C) Archaea: *P. furiosus*-EC [ PDB 9CRO (3)]. (D) Eukaryotic PolII: *S. cerevisiae* PolII-EC [PDB 9RYB (4)]. (E) Denaturing polyacrylamide gel showing results of single nucleotide extension assays for *Eco*RNAP in the presence of CHAPSO. (F) Denaturing polyacrylamide gel showing results of single nucleotide extension assays for *Mtb*RNAP in the presence of FC8F.

**Figure S3.**
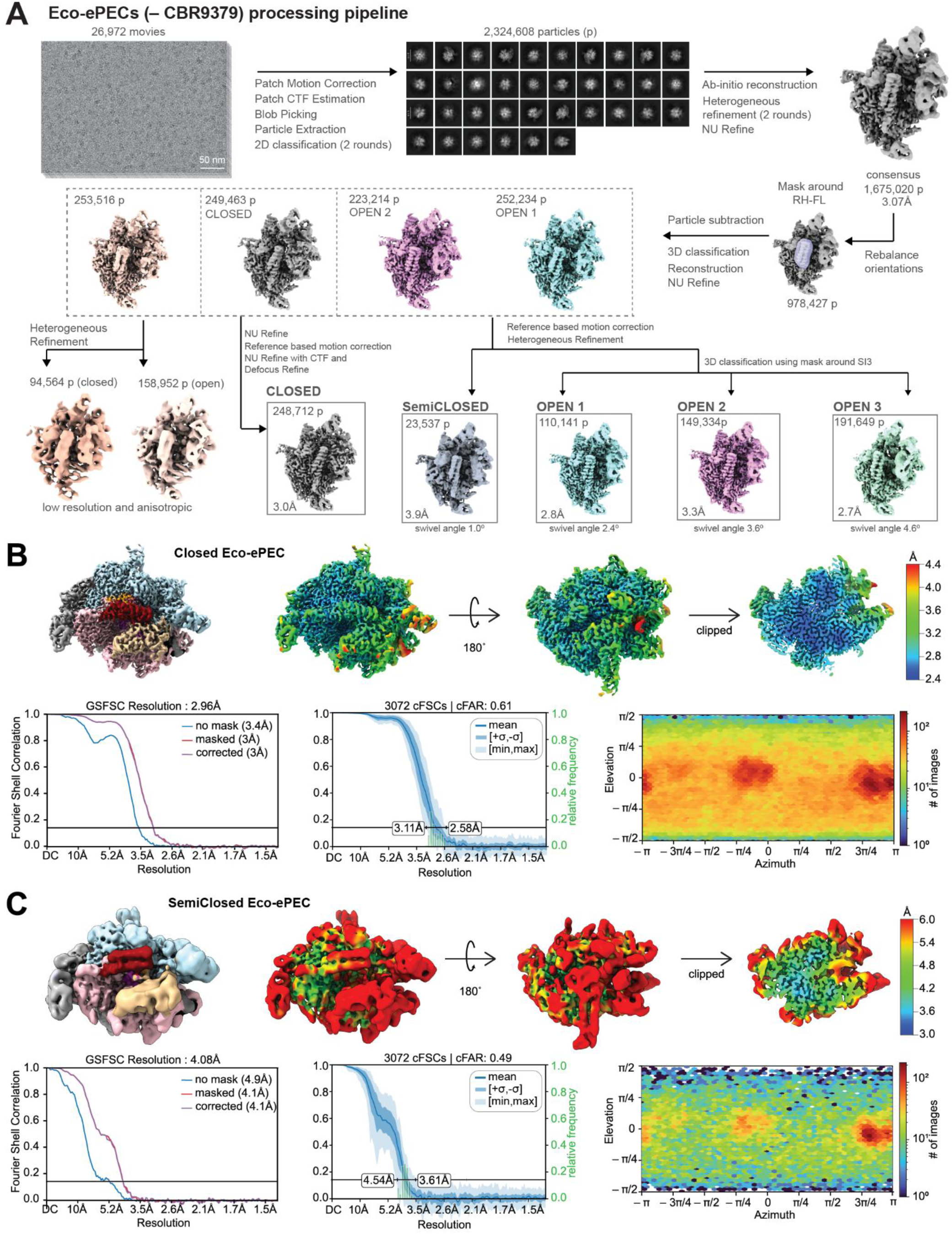
Cryo-EM data processing of *Eco*-ePECs(-CBR9379), local resolution, FSC, and particle orientation of Closed and SemiClosed *Eco*-ePECs. (A) Initial processing of raw movies, motion correction, CTF estimation, particle selection, 2D classification, to a consensus map and particle stack. The consensus particle stack was subjected to masked 3D classification around the RH-FL domain and further processed through heterogeneous refinements, more 3D classifications and reference-based motion corrections. Final reconstructions, Closed, SemiClosed, Open^1-3^ *Eco*-ePECs are boxed with particle number and resolution, and swivel angles for respective models noted under the box. See methods section for details. (B-C) Top panel: Locally filtered maps, colored by subunits and structural features (left), or colored by local resolution (scale on the right), rotated 180° and clipped (5). Bottom left panel: FSC curves calculated without mask (blue), masked (red), and corrected (purple). Bottom middle panel: 3D-FSC plot calculated by cryoSPARC showing mean FSC curve and FSC curves +/− 1 standard deviation. Bottom right panel: Heat map of particle orientation assignments. (B) Closed *Eco*-ePEC. (C) SemiClosed *Eco*-ePEC.

**Figure S4.**
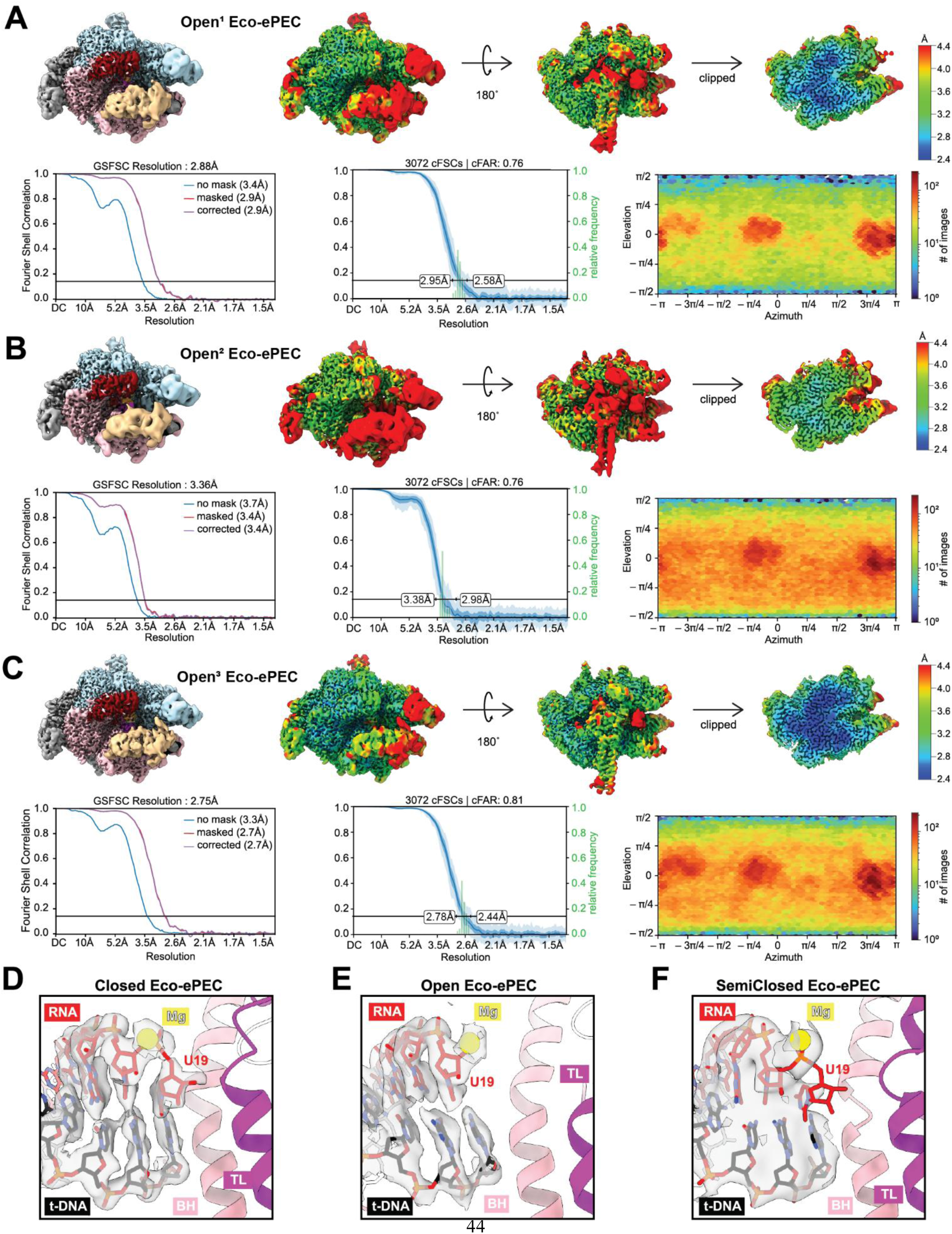
Local resolution, FSC, and particle orientation of Open^1-3^ *Eco*-ePECs; translocation states of *Eco*-ePECs. (A-C) Top panel: Locally filtered maps, colored by subunits and structural features (left), or colored by local resolution (scale on the right), rotated 180° and clipped (5). Bottom left panel: FSC curves calculated without mask (blue), masked (red), and corrected (purple). Bottome middle panel: 3D-FSC plot calculated by cryoSPARC showing mean FSC curve and FSC curves +/− 1 standard deviation. Bottom right panel: Heat map of particle orientation assignments. (A) Open^1^ *Eco*-ePEC. (B) Open^2^ *Eco*-ePEC. (C) Open^3^ *Eco*-ePEC. (D) Enlarged view of active-site Closed *Eco*-ePEC with map density (for t-DNA and RNA) shown, illustrating a pre-translocated state. U19 is labeled. (E) Enlarged view of active-site Open *Eco*-ePEC with map density (for t-DNA and RNA) shown, illustrating a half-translocated state. U19 is labeled. (F) Enlarged view of active-site SemiClosed *Eco*-ePEC with map density shown. U19 is labeled.

**Figure S5.**
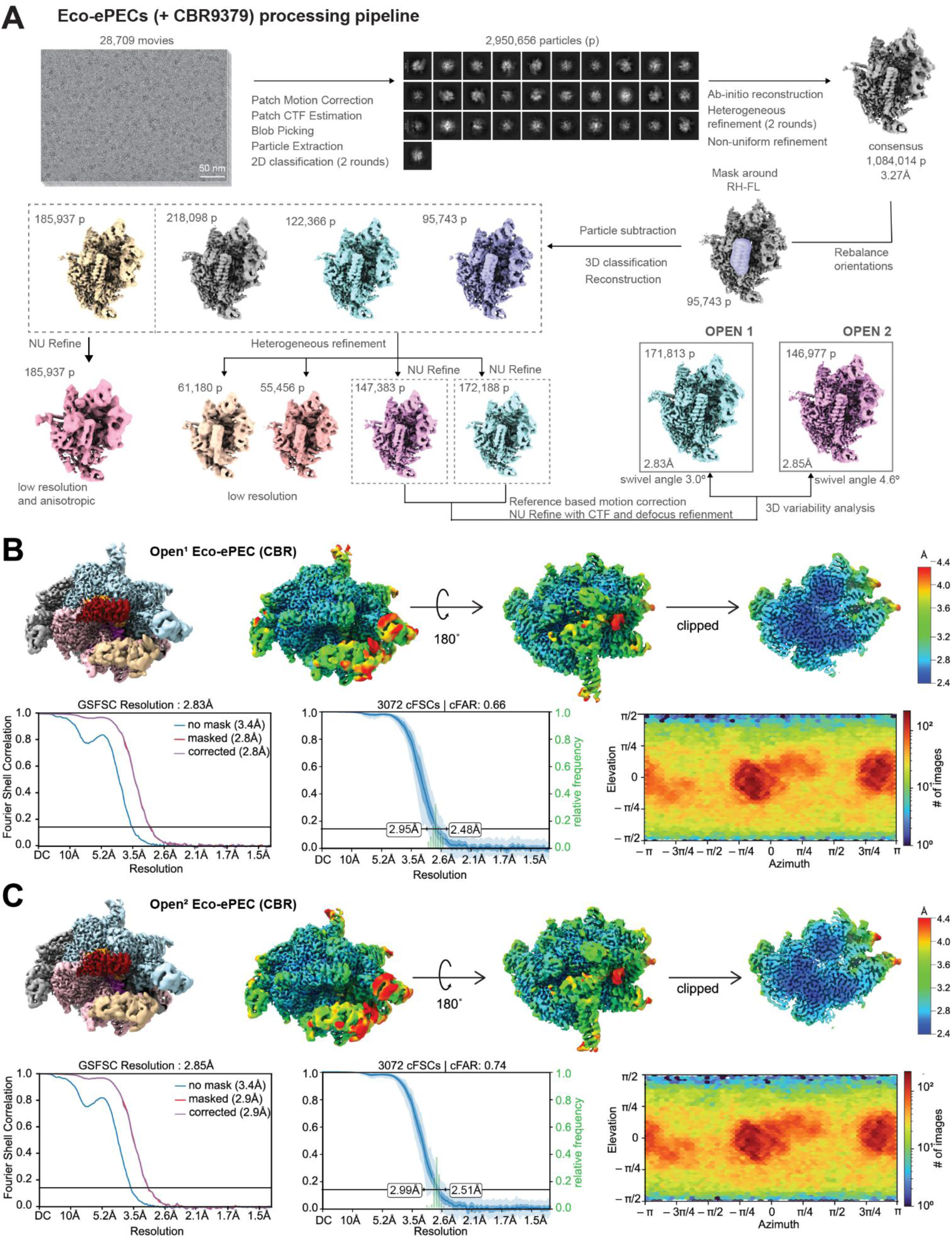
Cryo-EM data processing of *Eco*-ePECs(+CBR9379), Local resolution, FSC, and particle orientation of Open *Eco*-ePECs (CBR). (A) Initial processing of raw movies, motion correction, CTF estimation, particle selection, 2D classification, to a consensus map and particle stack. Consensus particle stack was subjected to masked 3D classification around the RH-FL domain and further processed through heterogeneous refinements, more 3D classifications and reference-based motion corrections. Final reconstructions Open^1^ *Eco*-ePEC(CBR9379) and Open^2^ *Eco*-ePEC(CBR9379) are boxed with particle number and resolution, and swivel angles for respective models noted under the box. See methods section for details. (B-C) Top panel: Locally filtered maps, colored by subunits and structural features (left), or colored by local resolution (scale on the right), rotated 180° and clipped (5). Bottom left panel: FSC curves calculated without mask (blue), masked (red), and corrected (purple). Bottom middle panel: 3D-FSC plot calculated by cryoSPARC showing mean FSC curve and FSC curves +/− 1 standard deviation. Bottom right panel: Heat map of particle orientation assignments. (B) Open^1^ *Eco*-ePEC(CBR9379). (C) Open^2^ *Eco*-ePEC(CBR9379).

**Figure S6.**
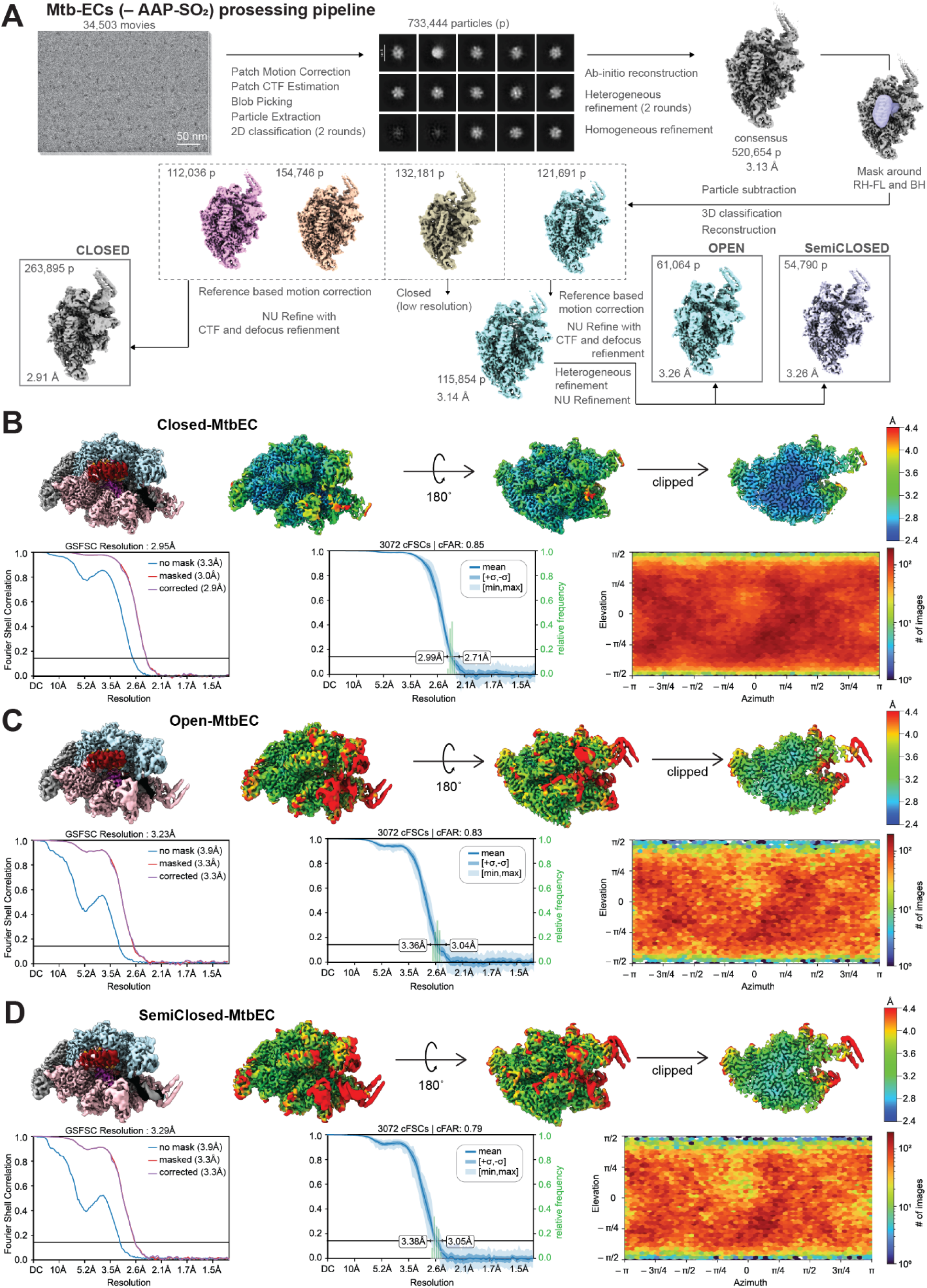
Cryo-EM data processing of *Mtb*-ECs(-AAP-SO_2_), Local resolution, FSC, and particle orientation of *Mtb*-ECs. (A) Initial processing of raw movies, motion correction, CTF estimation, particle selection, 2D classification, to a consensus map and particle stack. The consensus particle stack was subjected to masked 3D classification around the RH-FL domain and further processed through heterogeneous refinements, more 3D classifications and reference-based motion corrections. Final reconstructions, Closed, Open, SemiClosed *Mtb*-ECs are boxed with particle number and resolution. See methods section for details. (B-D) Top panel: Locally filtered maps, colored by subunits and structural features (left), or colored by local resolution (scale on the right), rotated 180° and clipped (5). Bottom left panel: FSC curves calculated without mask (blue), masked (red), and corrected (purple). Bottom middle panel: 3D-FSC plot calculated by cryoSPARC showing mean FSC curve and FSC curves +/− 1 standard deviation. Bottom right panel: Heat map of particle orientation assignments. (B) Closed *Mtb*-EC. (C) Open *Mtb*-EC. (D) SemiClosed *Mtb*-EC.

**Figure S7.**
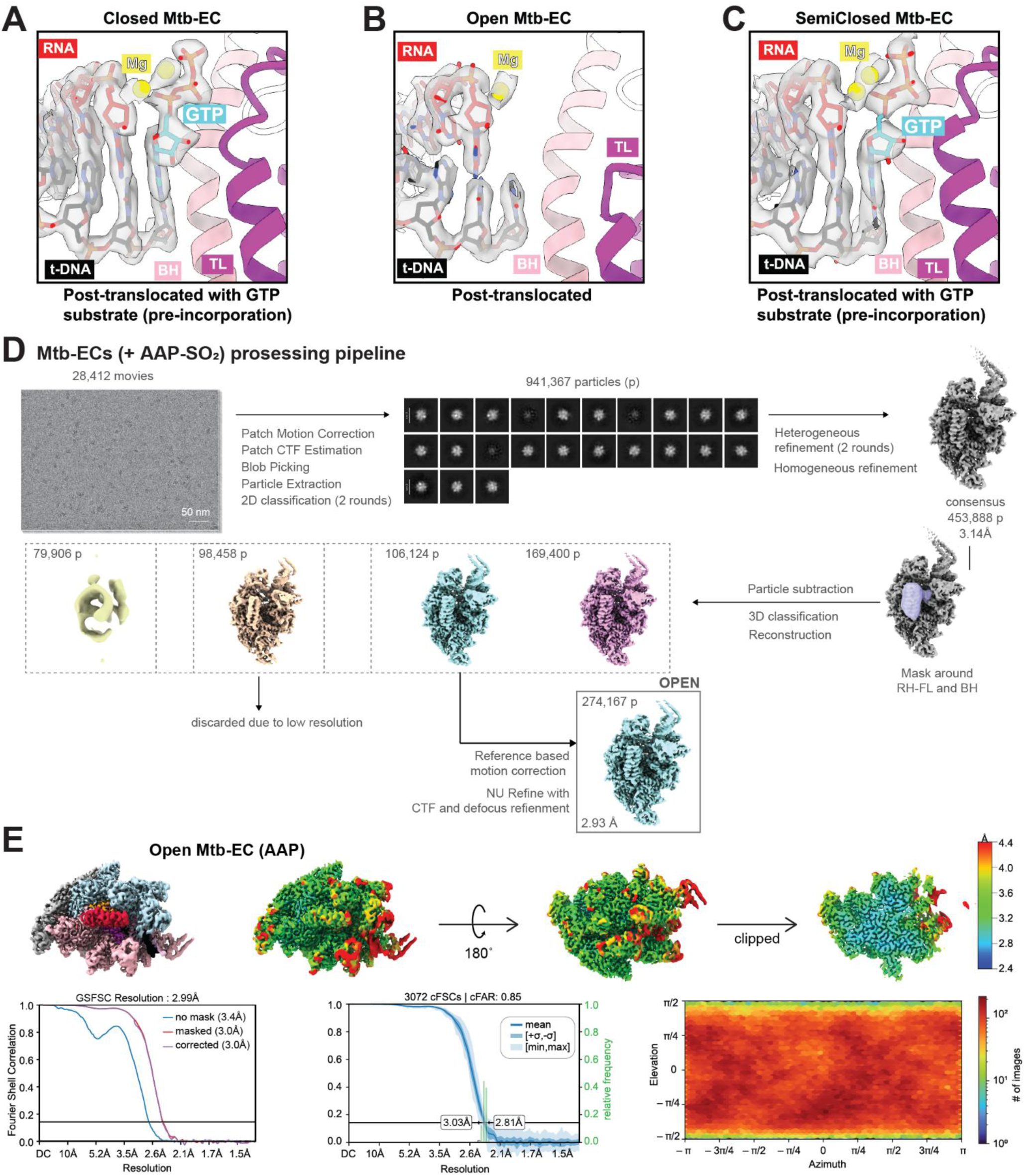
Translocations states of *Mtb*-ECs, Cryo-EM data processing pipeline of *Mtb*-ECs(+AAP-SO_2_), Local resolution, FSC, and particle orientation of Open *Mtb-*EC(AAP-SO_2_). (A) Enlarged view of active-site Closed *Mtb*-EC with map density (for t-DNA and RNA) shown, illustrating a pre-incorporation or substrate loading state of the NAC. Incoming GTP is shown in cyan. (A) Enlarged view of active-site Open *Mtb*-EC with map density (for t-DNA and RNA) shown, illustrating a post-translocated state. (C) Enlarged view of active-site SemiClosed *Eco*-ePEC with map density shown, illustrating a pre-incorporation or substrate loading state of the NAC. Incoming GTP is shown in cyan. (D) Initial processing of raw movies, motion correction, CTF estimation, particle selection, 2D classification, to a consensus map and particle stack. The consensus particle stack was subjected to masked 3D classification around the RH-FL domain and further processed through heterogeneous refinements, more 3D classifications and reference-based motion corrections. The final reconstruction, Open *Mtb*-ECs, is boxed with particle number and resolution. See methods section for details. (E) Top panel: Locally filtered maps, colored by subunits and structural features (left), or colored by local resolution (scale on the right), rotated 180° and clipped (5). Bottom left panel: FSC curves calculated without mask (blue), masked (red), and corrected (purple). Bottom middle panel: 3D-FSC plot calculated by cryoSPARC showing mean FSC curve and FSC curves +/− 1 standard deviation. Bottom right panel: Heat map of particle orientation assignments.

**Table S1.** Cryo-EM data collection, refinement and validation statistics for Eco-PECs.

|  | *Eco-ePEC* |  |  |  |  | *Eco-ePEC*<br>(CBR9379) |  |
| --- | --- | --- | --- | --- | --- | --- | --- |
| ***Data collection and processing*** |  |  |  |  |  |  |  |
| Magnification | 105000 |  |  |  |  | 105000 |  |
| Voltage (kV) | 300 |  |  |  |  | 300 |  |
| Electron exposure (e-/Å<sup>2</sup>) | 50 |  |  |  |  | 52.5 |  |
| Defocus range (μm) | 0.5 μm to 2.0 μm |  |  |  |  | 0.5 μm to 2.0 μm |  |
| Pixel size (Å) | 0.676 |  |  |  |  | 0.676 |  |
| Symmetry imposed | C1 |  |  |  |  | C1 |  |
| Initial particle images (no.) | 2,324,608 |  |  |  |  | 2,950,656 |  |
| Final (consensus) particle images (no.) | 1,675020 |  |  |  |  | 1,084,014 |  |
| Map (consensus) resolution (Å)<br>FSC threshold = 0.143 | 3.07 |  |  |  |  | 3.27 |  |
| ***Refinement*** |  |  |  |  |  |  |  |
| Complex | Closed | Open<sup>1</sup> | Open<sup>2</sup> | Open<sup>3</sup> | SemiClosed | Open<sup>1</sup> | Open<sup>2</sup> |
| PDB ID | 10MA | 10MB | 10MC | 10MD | 10ME | 10MF | 10MG |
| EMDB ID | EMD-75279 | EMD-75280 | EMD-75281 | EMD-75282 | EMD-75283 | EMD-75284 | EMD-75285 |
| *Initial model used (PDB code)* | 8EH8 | 8EHI | 8EHI | 8EHI | 8EHA | 8EHI | 8EHI |
| Model resolution (Å)<br>FSC threshold = 0.5 | 2.9 | 2.9 | 3.3 | 2.7 | 4.1 | 2.8 | 2.8 |
| Model composition |  |  |  |  |  |  |  |
| Non-hydrogen atoms | 26880 | 26776 | 26776 | 26776 | 26251 | 26979 | 26918 |
| Protein residues | 3190 | 3168 | 3168 | 3168 | 3190 | 3168 | 3168 |
| Nucleic acid residues | 64 | 63 | 63 | 63 | 64 | 63 | 63 |
| Ligands | 4 | 4 | 4 | 4 | 4 | 5 | 5 |
| B factors (Å<sup>2</sup>) |  |  |  |  |  |  |  |
| Protein | 52.67 | 60.39 | 174.58 | 75.34 | 243.23 | 68.16 | 67.20 |
| Nucleic acids | 105.99 | 87.64 | 228.61 | 121.29 | 259.92 | 135.86 | 105.69 |
| Ligand | 48.45 | 41.74 | 151.44 | 57.00 | 228.32 | 47.67 | 44.35 |
| R.m.s. deviations |  |  |  |  |  |  |  |
| Bond lengths (Å) | 0.006 | 0.002 | 0.003 | 0.003 | 0.004 | 0.005 | 0.002 |
| Bond angles (°) | 0.894 | 0.558 | 0.598 | 0.576 | 0.723 | 0.692 | 0.579 |
| Validation |  |  |  |  |  |  |  |
| MolProbity score | 1.97 | 1.53 | 1.55 | 1.43 | 2.18 | 1.84 | 1.62 |
| Clashscore | 4.67 | 2.81 | 3.39 | 2.50 | 7.70 | 3.31 | 2.98 |
| Poor rotamers (%) | 4.84 | 3.12 | 2.93 | 3.04 | 4.66 | 5.08 | 3.53 |
| Ramachandran plot |  |  |  |  |  |  |  |
| Favored (%) | 96.66 | 97.62 | 97.71 | 97.87 | 96.28 | 96.92 | 97.40 |
| Allowed (%) | 3.34 | 2.38 | 2.29 | 2.13 | 3.72 | 3.08 | 2.60 |
| Disallowed (%) | 0.00 | 0.00 | 0.00 | 0.00 | 0.00 | 0.00 | 0.00 |

**Table S2.** Cryo-EM data collection, refinement and validation statistics for Mtb-ECs.

|  | *Mtb*-EC |  |  | *Mtb*-EC (AAP-SO<sub>2</sub>) |
| --- | --- | --- | --- | --- |
| ***Data collection and processing*** |  |  |  |  |
| Magnification | 105,000 |  |  | 105,000 |
| Voltage (kV) | 300 |  |  | 300 |
| Electron exposure (e-/Å<sup>2</sup>) | 50.7 |  |  | 50.7 |
| Defocus range (µm) | 0.7 µm to 2.0 µm |  |  | 0.7 µm to 2.0 µm |
| Pixel size (Å) | 0.847 |  |  | 0.847 |
| Symmetry imposed | C1 |  |  | C1 |
| Initial particle images (no.) | 733,444 |  |  | 941,367 |
| Final (consensus) particle images (no.) | 520,654 |  |  | 454,888 |
| Map (consensus) resolution (Å)<br>FSC threshold = 0.143 | 3.13 |  |  | 3.14 |
| ***Refinement*** |  |  |  |  |
| Complex | Closed | Open | SemiClosed | Open |
| PDB ID | 10MI | 10MJ | 10MK | 10ML |
| EMDB ID | EMD-75286 | EMD-75287 | EMD-75288 | EMD-75289 |
| *Initial model used (PDB code)* | 8e95 | 8e95 | 8e95 | 8e95, 9MRQ |
| Model resolution (Å)<br>FSC threshold = 0.5 | 2.9 | 3.3 | 3.3 | 3.0 |
| Model composition |  |  |  |  |
| Non-hydrogen atoms | 24989 | 24201 | 24989 | 24631 |
| Protein residues | 2918 | 2910 | 2918 | 2909 |
| Nucleic acid residues | 70 | 61 | 70 | 61 |
| Ligands | 5 | 3 | 5 | 6 |
| B factors (Å<sup>2</sup>) |  |  |  |  |
| Protein | 42.58 | 59.04 | 63.83 | 6.44 |
| Nucleic acids | 64.08 | 137.23 | 121.77 | 13.73 |
| Ligand | 17.73 | 84.69 | 43.32 | 10.93 |
| R.m.s. deviations |  |  |  |  |
| Bond lengths (Å) | 0.003 (0) | 0.002 (0) | 0.004 (0) | 0.004 (0) |
| Bond angles (°) | 0.577 (0) | 0.521 (0) | 0.567 (0) | 0.822 (0) |
| Validation |  |  |  |  |
| MolProbity score | 1.59 | 1.78 | 1.57 | 1.64 |
| Clashscore | 2.73 | 3.18 | 2.46 | 2.79 |
| Poor rotamers (%) | 4.79 | 4.34 | 3.48 | 3.73 |
| Ramachandran plot |  |  |  |  |
| Favored (%) | 97.97 | 96.92 | 97.31 | 97.73 |
| Allowed (%) | 2.03 | 3.08 | 2.69 | 2.77 |
| Disallowed (%) | 0.00 | 0.00 | 0.00 | 0.00 |

